# Divergent *Rickettsia* species exhibit distinct mechanisms of actin-based motility

**DOI:** 10.1101/2025.08.14.669974

**Authors:** Meghan C. Bacher, Julie E. Choe, Jiawen Jiang, Joanna M. Idrovo, Joshua T. Del Mundo, Michal Hammel, Matthew D. Welch

## Abstract

Many *Rickettsia* species undergo actin-based motility to promote cell-cell spread during infection. Rickettsial genomes encode two motility effectors, RickA and Sca2. In the spotted fever group species *R. parkeri*, RickA acts early in infection by activating the host Arp2/3 complex; Sca2 acts later by mimicking the structure and function of eukaryotic formins. The function of RickA and Sca2 orthologs in the distantly related species *R. bellii* was unclear. We report that the *R. bellii* Sca2 ortholog, Sca2/6, nucleates and elongates actin with a flexible structure and an unusual actin monomer-binding motif in a mechanism distinct from formins or other microbial actin nucleators. *R. bellii* motility occurs only later in infection and is solely correlated with Sca2/6 localization. Compared with *R. parkeri*, *R. bellii* motility is slow and meandering, and generates distinctly organized actin tails, reflecting differences in Sca2 ortholog mechanism and localization. The evolutionary flexibility in the mechanism and regulation of rickettsial actin-based motility suggests similar adaptability for other microbes.

## Introduction

Intracellular bacteria have evolved numerous ways to co-opt the host cell actin cytoskeleton for processes including invasion, innate immune avoidance, and cell-cell spread (Colonne et al., 2016; Lamason and Welch, 2017; Stradal and Schelhaas, 2018). Actin-based motility is one mechanism through which actin contributes to microbial cell-cell spread. This process uses the force derived from the nucleation and elongation of actin filaments at the bacterial surface to drive movement through the host cell cytoplasm, resulting in the assembly of actin comet tails (Lamason and Welch, 2017; Dowd et al., 2021). Motility positions bacteria near the host cell plasma membrane to enable them to enter membrane protrusions that can be engulfed into neighboring cells (Lamason and Welch, 2017; Dowd et al., 2021).

The mechanism of actin-based motility has been studied for a variety of pathogens including *Rickettsia* species (Lamason and Welch, 2017; Alqassim, 2022). *Rickettsia* are Gram-negative obligate intracellular alphaproteobacteria that typically reside in arthropods such as ticks, fleas, and mites (Voss and Rahman, 2021). The *Rickettsia* genus is divided into 5 distinct groups clustered by molecular phylogenomics and disease features (Karkouri et al., 2022; Rymaszewska and Piotrowski, 2024). These groups include the spotted fever groups (SFG I/II) and typhus group (TG), which contain pathogens that cause spotted fever and typhus in humans. They also include the canadensis group (CG) and bellii group (BG), which contain species considered to be non-pathogenic tick endosymbionts. Actin-based motility has been observed for divergent species within most of the groups including the SFG I species, *R. parkeri*, and the BG species, *R. bellii* (Teysseire et al., 1992; Heinzen et al., 1993; Gouin et al., 1999; Van Kirk et al., 2000; Haglund et al., 2010; Pan et al., 2022).

*Rickettsia* species are distinctive amongst microbes that undergo actin-based motility in that their genomes often encode two different motility effectors, RickA and Sca2 (Kleba et al., 2010; Haglund et al., 2010; Reed et al., 2014; Lehman et al., 2024). Studies with the SFG I species, *R. parkeri*, harboring transposon mutants in *rickA* or *sca2* demonstrated that RickA and Sca2 function independently to drive two temporal phases of motility (Reed et al., 2014; Harris et al., 2018). RickA-motility occurs at early time points of infection (< 2 h post-infection (hpi)), is slower, and proceeds in meandering trajectories, resulting in actin tails that are short and curved (Reed et al., 2014; Jeng et al., 2004). Sca2-motility occurs later (> 5 hpi), is faster, and proceeds in linear trajectories, resulting in actin tails that are long and straight (Reed et al., 2014). Mutations in *rickA* and *sca2* lead to the formation of smaller plaques during infection, indicating both RickA-motility and Sca2-motility contribute to cell-cell spread (Kleba et al., 2010; Reed et al., 2014; Tran et al., 2025). However, the modes of spread differ in key parameters.

RickA-spread results from the formation of longer and more dynamic plasma membrane protrusions (Tran et al., 2025), whereas Sca2-spread results from the formation of shorter and less dynamic protrusions (Lamason et al., 2016; Tran et al., 2025). In a mouse model of infection, RickA is important for eschar formation in the skin, whereas Sca2 is important for dissemination to internal organs (Burke et al., 2021; Tran et al., 2025).

In addition to differences in their roles in cell-cell spread and pathogenicity, RickA and Sca2 from SFG I species nucleate actin filaments through different biochemical mechanisms. RickA is secreted by bacteria through an unknown mechanism and acts as a nucleation-promoting factor (NPF) that activates the host Arp2/3 complex through conserved C-terminal WH2, central, and acidic (WCA) motifs (Gouin et al., 2004; Jeng et al., 2004). Activated Arp2/3 complex nucleates a new actin filament from the side of an existing filament in a Y-branched configuration (Gautreau et al., 2022). Sca2 is a type V secretion system (T5SS) or autotransporter protein, a family of proteins that consists of an N-terminal signal sequence that enables passage through the bacterial inner membrane, a C-terminal translocation domain consisting of a β-barrel structure that embeds in the outer membrane, and a central passenger domain that threads through the translocation domain and is displayed on the bacterial surface (Fan et al., 2016). The Sca2 passenger domain from SFG I species nucleates and processively elongates actin filaments in a profilin-dependent manner, similar to eukaryotic formin proteins (Haglund et al., 2010; Madasu et al., 2013; Alqassim et al., 2019). Within the passenger domain of SFG I Sca2, the middle domain binds to monomeric globular (G) actin, and the N-terminal domain (NTD) folds into a ring structure resembling a formin-like core that cooperatively binds two actin subunits with high affinity in conjunction with the C-terminal repeat domain (CRD) (Alqassim et al., 2019; Carman et al., 2023). Interestingly, the passenger domain of the Sca2 ortholog from the divergent BG species, *R. bellii*, differs dramatically in sequence from SFG I Sca2 (Haglund et al., 2010). The single gene encoding the *R. bellii* Sca2 ortholog may have undergone a tandem duplication during rickettsial evolution, yielding adjacent *sca2* and *sca6* genes (Blanc et al., 2005; Ngwamidiba et al., 2005). Both genes are intact in the SFG II species, *R. akari*, whereas one gene is degraded in SFG I species, and one or both are degraded in TG species (Blanc et al., 2005; Ngwamidiba et al., 2005; Sears et al., 2012). For this reason, the *R. bellii* ortholog is referred to as Sca2/6 (Lehman et al., 2024). We do not yet know whether and how sequence or regulatory differences in RickA or Sca2 orthologs contribute to differences in the mechanism and properties of actin-based motility in diverse *Rickettsia* species.

To better understand the functions of actin-based motility proteins in divergent *Rickettsia* species, we compared the actin nucleation mechanisms, protein localization, and motility characteristics between the non-pathogenic tick endosymbiont *R. bellii* and the well-studied pathogenic SFG I species *R. parkeri*. We found that the Arp2/3-dependent actin nucleation mechanism of RickA is conserved. However, *R. bellii* Sca2/6 nucleates and elongates actin filaments by a mechanism that differs from the formin-like mechanism of SFG I Sca2.

Unexpectedly, in infected cells, *R. bellii* motility did not occur early in infection, was functionally independent of RickA and the Arp2/3 complex, and was tightly correlated with highly polar Sca2/6 localization. Differences in mechanism and localization between *R. bellii* Sca2/6 and *R. parkeri* Sca2 orthologs also correlated with differences in actin tail organization and actin-based motility. Together, these data indicate that there is considerable evolutionary flexibility in actin-based motility proteins and mechanisms within the *Rickettsia* genus.

## Results

### RickA is conserved while Sca2/6 has diverged in *R. bellii*

To better understand how RickA and Sca2 have diverged across *Rickettsia* species, we aligned the protein sequences from representative *Rickettsia* species from the SFG I/II, TG (Sca2 only), and BG groups, including *R. parkeri* strain Portsmouth and *R. bellii* strain RML 369-C. For RickA, the overall organization of sequence motifs is conserved (Figure 1A). Alignment of RickA protein sequences (Figure S1A) showed that the C-terminal WCA region exhibits 27% sequence identity and 56% similarity overall. SFG I and BG species contain a single WH2 motif, whereas SFG II species contain two WH2 motifs (Figure 1A and S1A). This high degree of conservation suggests that the biochemical function and activities of RickA orthologs are conserved between species.

**FIGURE 1.**
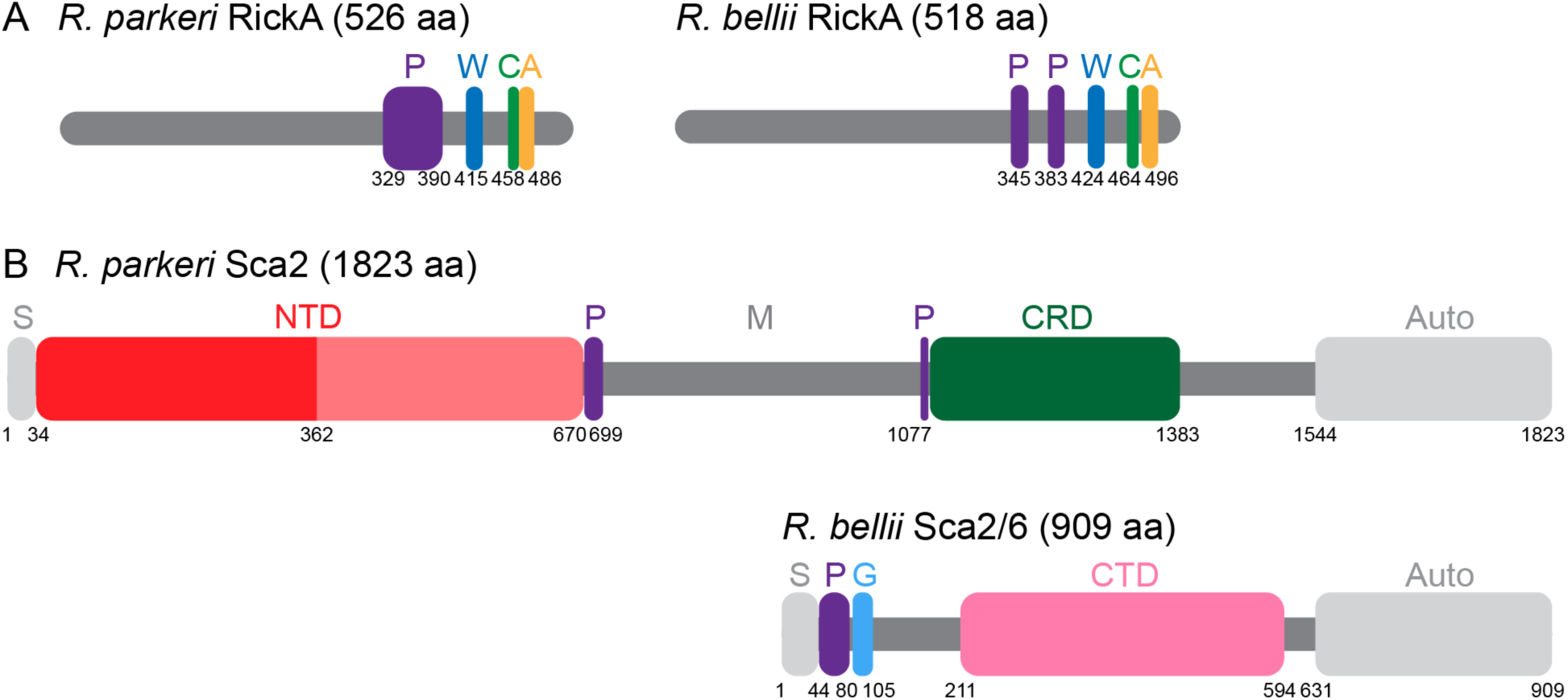
RickA domain organization is conserved, whereas Sca2 ortholog domain organization is more variable. (A) RickA from *R. parkeri* and *R. bellii* share a common domain architecture. (B) Sca2 from *R. parkeri* and Sca2/6 *R. bellii* proteins differ considerably in their domain architecture. Abbreviations in (A) and (B): P = poly-proline or proline-rich domain, W = WASP homology-2 (WH2) motif, C = central motif, A = acidic motif, S = signal sequence, NTD = N-terminal domain (the darker half denotes the location of helical N-terminal repeats), M = middle domain, G = G-actin binding domain, CRD = C-terminal repeat domain, CTD = C-terminal domain, Auto = autotransporter domain.

In contrast, Sca2 orthologs from divergent species differ considerably in the organization of sequence motifs in the passenger domain (Figure 1B). In comparison with Sca2 from the SFG I species *R. parkeri* (1823 amino acids), Sca2/6 from *R. bellii* is half as long (909 amino acids) and lacks the N-terminal repeats within the NTD and CRD domains that were previously shown to form a partially occluded formin-like core (Figure 1B, S1B) (Haglund et al., 2010; Madasu et al., 2013; Alqassim et al., 2019; Carman et al., 2023). The passenger domains of Sca2 orthologs only exhibit 11% percent identity and 33% similarity (Figure S1B). Using a combination of AlphaFold 3 (Abramson et al., 2024) and InterPro (Blum et al., 2024) protein domain predictions, we annotated the passenger domain of *R. bellii* Sca2/6 as having a signal sequence (S), a poly-proline region (P), putative a G-actin binding motif (G) previously annotated as a WH2 motif (Haglund et al., 2010), and a C-terminal domain (CTD) (Figure 1B, S1B). As described for Sca2 from the SFG I species *R. conorii*, the previously annotated WH2 motifs in *R. bellii* Sca2/6 (Haglund et al., 2010) are predicted to have different secondary structures than canonical WH2 domains (Ducka et al., 2010; Dominguez, 2016). Nevertheless, AlphaFold 3 structure predictions suggested that an α-helix within the previously annotated WH2 motif of *R. bellii* Sca2/6 binds to one G-actin molecule at its barbed end (pLDDT > 70, Figure S2). This updated analysis confirms that the passenger domain of the Sca2 orthologs differs substantially between *R. bellii* and the SFG I species *R. parkeri*, suggesting potential differences in biochemical activities and mechanisms of action.

We also sought to confirm that *R. bellii* expresses RickA and Sca2/6. Both proteins were detected in *R. bellii* bacterial lysates by immunoblotting with anti-RickA or anti-Sca2/6 antibodies (Figure S3). Thus, *R. bellii* expresses both actin-based motility effector proteins.

### *R. bellii* RickA activates the host Arp2/3 complex *in vitro*

RickA proteins from the SFG I species, *R. rickettsii* (Jeng et al., 2004) and *R. conorii* (Gouin et al., 2004), were previously shown to function as NPFs and activate the host Arp2/3 complex. To test whether RickA from *R. bellii* also possesses NPF activity, we expressed and purified glutathione S-transferase (GST) tagged RickA (GST-RickA) and performed pyrene actin polymerization assays (Figure S4). Purified GST-RickA or Arp2/3 complex individually did not accelerate actin polymerization compared with actin alone. However, when combined, GST-RickA activated the Arp2/3 complex to nucleate actin in a concentration-dependent manner. These data confirm that *R. bellii* RickA has NPF activity.

### *R. bellii* Sca2/6 nucleates and elongates actin filaments *in vitro*

Our sequence comparisons of Sca2 orthologs from *R. parkeri* and *R. bellii* suggested that the biochemical activity of *R. bellii* Sca2/6 may differ considerably from that of *R. parkeri* Sca2, which nucleates actin assembly and caps actin filaments in the absence of profilin (Haglund et al., 2010; Madasu et al., 2013). To determine whether *R. bellii* Sca2/6 nucleates actin assembly, we expressed and purified the maltose-binding protein tagged passenger domain of Sca2/6 (MBP-Sca2/6; excluding the signal sequence and autotransporter domains) and performed pyrene actin polymerization assays. Purified MBP-Sca2/6 alone nucleated actin assembly in a concentration-dependent manner (Figure 2A). Cleavage of the MBP tag did not significantly impact actin nucleation activity (Figure S5A). For comparison, *R. parkeri* GST-Sca2 alone also nucleated actin filament assembly, but fluorescence plateaued at a lower value due to its barbed end capping activity in the absence of profilin (Haglund et al., 2010) (Figure 2A). Thus, *R. bellii* Sca2/6 nucleates actin assembly without capping the barbed ends of actin filaments.

**FIGURE 2.**
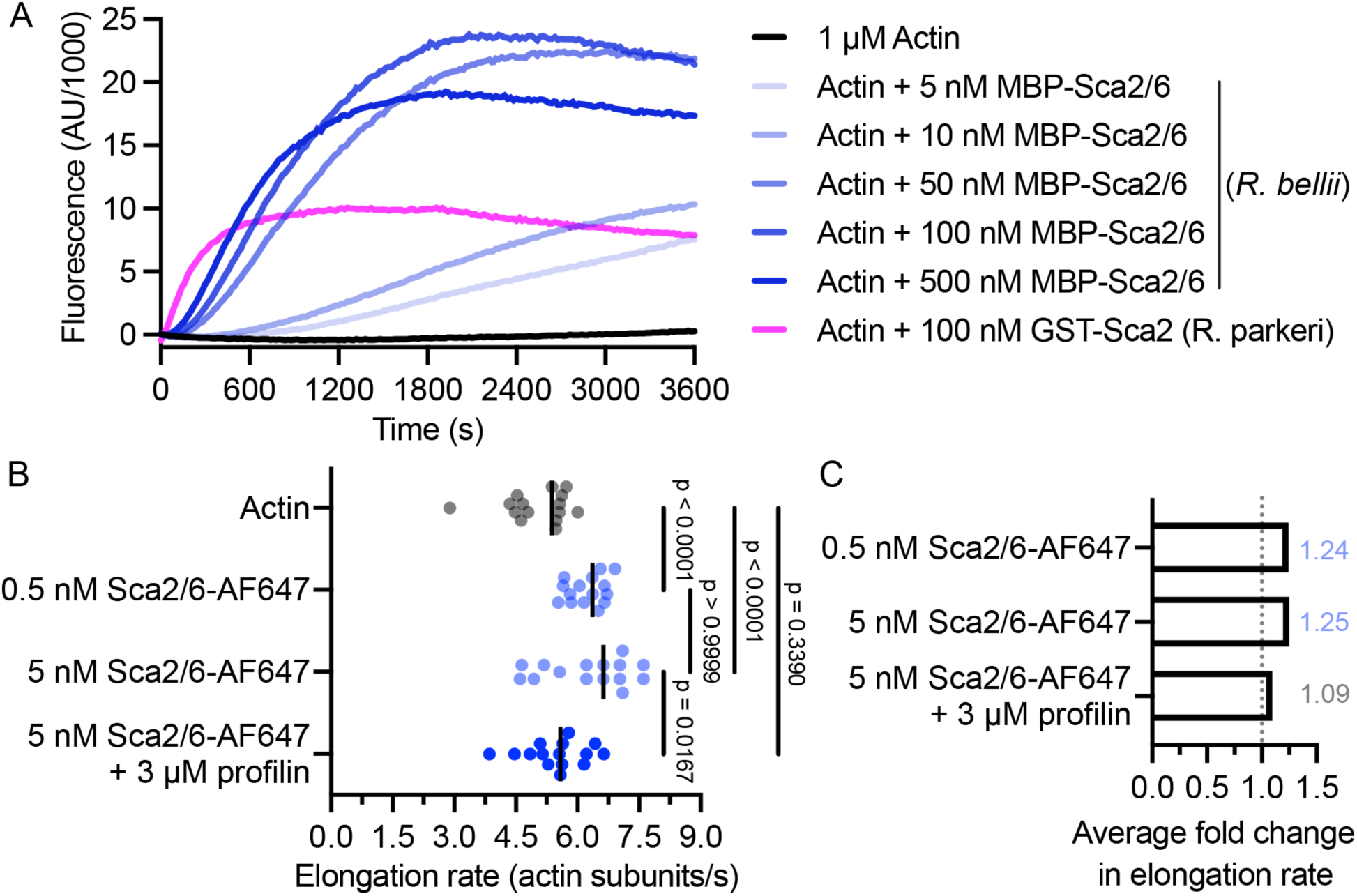
*R. bellii* Sca2/6 nucleates actin filaments and promotes filament elongation. (A) Kinetics of assembly of 1 µM actin (10% pyrene-labeled) with the indicated concentrations of the *R. bellii* MBP-Sca2/6 or *R. parkeri* GST-Sca2 passenger domains. (B) Quantification of actin filament barbed-end elongation rates in TIRF assays performed with 1 µM actin (33% rhodamine-labeled) without or with the indicated concentrations of *R. bellii* Sca2/6-AF647 and 3 µM profilin. Vertical black lines indicate median elongation rate. N = 3 independent experiments per condition, n = 5 filaments per experiment. Significance was determined using a one-way ANOVA with a Tukey’s multiple comparison post-test. (D) Average fold change in barbed end actin filament elongation rates for the indicated concentrations of Sca2/6-AF647 and profilin relative to actin alone (grey dotted line).

*R. parkeri* Sca2 processively associates with barbed ends and enables filament elongation in the presence of profilin (Haglund et al., 2010; Madasu et al., 2013), like eukaryotic formins (Breitsprecher and Goode, 2013; Valencia and Quinlan, 2021). For comparison, we sought to assess whether *R. bellii* Sca2/6 stably associates with actin filaments and/or affects the rate of filament elongation using total internal reflection fluorescence (TIRF) microscopy. We fluorescently labeled the C-terminus of *R. bellii* Sca2/6 with Alexa Fluor 647 (AF647) and cleaved the MBP-tag to create Sca2/6-AF647. Sca2/6-AF647 and MBP-Sca2/6 activity were similar in pyrene actin assays (Figure S5). Polymerization of 1 µM actin was initiated in the absence or presence of Sca2/6-AF647, and the growth of actin filaments was visualized with TIRF microscopy. We did not observe stable association of Sca2/6 puncta with polymerizing actin filaments over a range of Sca2/6-AF647 concentrations (Movie S1). However, transient interactions between Sca2/6-AF647 puncta and the sides or ends of growing filaments were sometimes spotted. The rate of barbed end growth was modestly (∼1.25-fold) but statistically significantly increased in the presence of Sca2/6-AF647 when compared with actin alone (Figure 2B and C). The addition of 3 µM profilin to Sca2/6-AF647 diminished the elongation rate to that of actin alone (Figure 2B and C). Together, these data show that *R. bellii* Sca2/6 nucleates actin and modestly enhances actin elongation independent of profilin and without detectable processive association with filament ends.

### *R. bellii* Sca2/6 is monomeric and structurally flexible in solution, binds G-actin, and requires G-actin binding for nucleation activity

To determine which domains of *R. bellii* Sca2/6 contribute to its actin nucleation activity, we expressed and purified a series of truncation mutants including: MBP-Sca2/6-ΔpolyP (missing the poly-proline region), MBP-Sca2/6-NTD (missing the CTD), or MBP-Sca2/6-CTD (missing the N-terminus) (Figure 3A). We performed pyrene actin assembly assays with 70 nM of each protein and calculated the maximum slope of the polymerization curve to compare their actin nucleation activities. MBP-Sca2/6-ΔpolyP exhibited activity that was not significantly different from full-length Sca2/6 (Figure 3B). In contrast, the activities of MBP-Sca2/6-NTD or MBP-Sca2/6-CTD were significantly reduced in comparison with full-length Sca2/6 and were comparable to actin alone. These data suggest that sequences outside the poly-proline motif are sufficient for full nucleation activity, and that sequences in both the N-terminus and CTD are necessary for activity.

**FIGURE 3.**
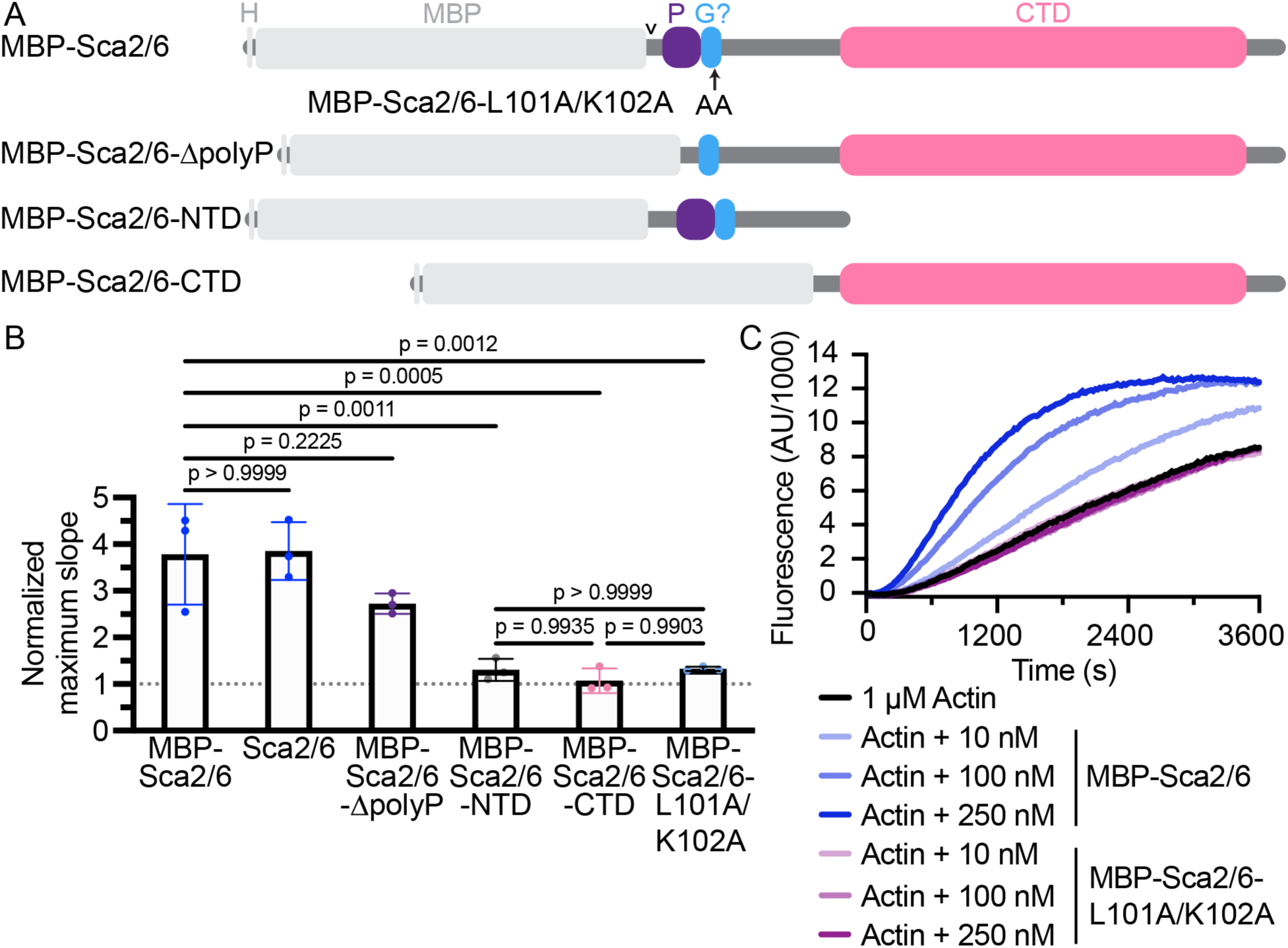
Actin nucleation activity of *R. bellli* Sca2/6 truncations and mutants. (A) Diagram of truncated and mutated derivatives of *R. bellii* Sca2/6. H = 6x-His tag, MBP = maltose-binding protein tag, P = poly-proline domain, G = G-actin binding domain, CTD = C-terminal domain, arrowhead = TEV protease cleavage site. The MBP-Sca2/6-L101A/K102A protein contains two point mutations in the G-actin binding motif. (B) Calculated maximum slope from pyrene-actin polymerization assays, normalized to actin alone (grey dotted line), for the indicated *R. bellii* Sca2/6 variants at 70 nM with average values noted in the bars. N = 3 experiments for each protein, n = 2 replicates per experiment. Error bars indicate SD. Significance was determined using a one-way ANOVA with a Tukey’s multiple comparisons test. (C) Kinetics of assembly of 1 µM actin (10% pyrene-labeled) with the indicated concentrations of *R. bellii* MBP-Sca2/6 or MBP-Sca2/6-L101A/K102A.

We next tested the function of the AlphaFold 3-predicted G-actin binding motif, comprising amino acids 86-105. To this end, we expressed and purified a variant of Sca2/6 in which amino acids L101 and K102 within this motif were substituted with alanine (MBP-Sca2/6-L101A/K102A). These amino acid substitutions substantially reduced the confidence of AlphaFold 3 predictions for actin-binding (pLDDT < 50, Figure S2). To experimentally test whether this motif binds G-actin, we performed small angle X-ray scattering in line with size exclusion chromatography (SEC-SAXS) with MBP-Sca2/6 alone as well as MBP-Sca2/6 or MBP-Sca2/6-L101A/K102A in the presence of G-actin (and latrunculin A to prevent actin polymerization). The calculated molecular weight (MW) from SAXS (based on the volume of correlation V_c_) of MBP-Sca2/6 was ∼110 kDa (sequence MW, 110 kDa), consistent with Sca2/6 behaving as a monomer in solution. In the presence of G-actin, the calculated MW for MBP-Sca2/6 plus actin was ∼125 kDa, and for MBP-Sca2/6-L101A/K102A plus actin was ∼105 kDa. The normalized Kratky plot shows that all measured proteins adopt a partially unfolded conformation (Figure S6). We then calculated pair distance distribution functions (P(r)) for MBP-Sca2/6, MBP-Sca2/6 plus actin, and MBP-Sca2/6-L101A/K102A plus actin. The extended P(r) “tails” at r > 170 Å in the P(r) plot indicate that in all samples, the *R. bellii* Sca2/6 protein adopts an extended conformation (Figure 4A). The peak at ∼50 Å corresponds to the intra-domain distance of the globular N-terminal MBP-tag and the broad peak at ∼120 Å corresponds to the flexible Sca2/6 protein (Figure 4A). The P(r) function for MBP-Sca2/6 plus actin additionally had a broad shoulder between ∼70-140 Å that is absent in functions derived for both MBP-Sca2/6 alone and MBP-Sca2/6-L101A/K102A plus actin. Thus, this shoulder contains both the intra-domain distance of actin and inter-domain distances with actin (Figure 4A). However, the significant broadening of the shoulder suggests that the Sca2/6 or Sca2/6-actin region is not compact and most likely adopts a flexible conformation in solution. In further support of the location of the G-actin binding motif, we visualized AlphaFold 3-derived predicted aligned errors (PAE), which measure the confidence in the relative position of two residues within the predicted structure (Figure 4B). MBP-Sca2/6 plus actin showed confidence in interactions with actin of Sca2/6 amino acids 70-110 (which includes L101 and K102). The confidence was significantly weaker for MBP-Sca2/6-L101A/K102A (Figure 4B). These data suggest that Sca2/6 binds one molecule of G-actin, and Sca2/6-L101A/K102A exhibits substantially reduced binding.

**FIGURE 4.**
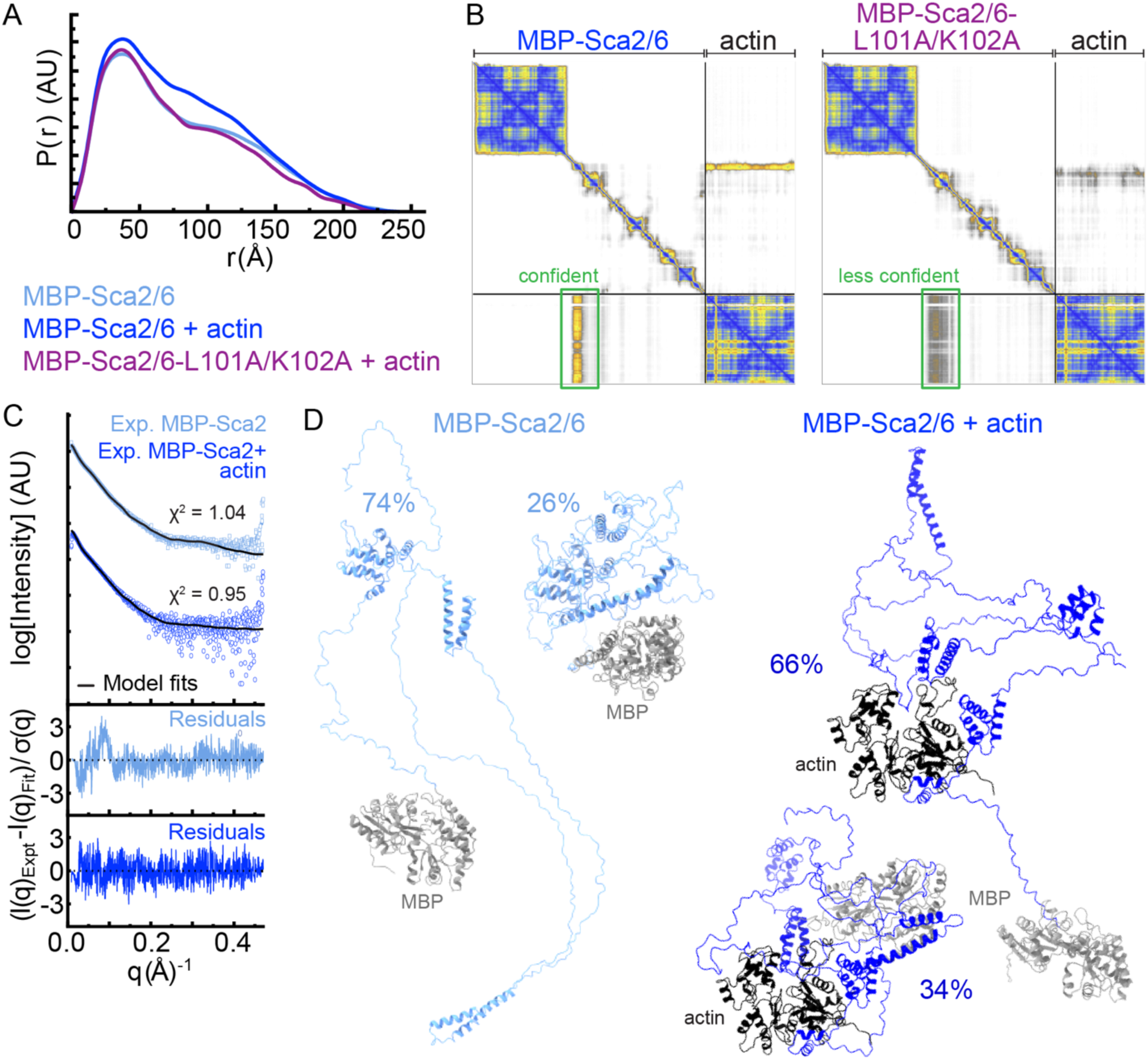
*R. bellii* Sca2/6 is a structurally flexible protein that binds one molecule of G-actin through an unusual G-actin binding motif in the N-terminus. (A) Pair distance distribution functions (P(r)) calculated from SAXS data for MBP-Sca2/6, MBP-Sca2/6 plus G-actin, and MBP-Sca2/6-L101A/K102A plus G-actin. Area under P(r) curves is proportional to the theoretical MW calculated from SAXS. (B) AlphaFold3 PAE matrices of MBP-Sca2/6 plus G-actin and MBP-Sca2/6-L101A/K102A plus G-actin. Green boxes highlight the clusters of higher or lower confidence values that indicate interactions between MBP-Sca2/6 or MBP-Sca2/6-L102A/K102A and G-actin. (C) Plot of SAXS experimental data (open circles and squares), BilboMD 2-state ensemble fits (black curves), and residuals. (D) BilboMD ensemble structures of MBP-Sca2/6 and MBP-Sca2/6 plus G-actin, labeled with the percentage of each structure in the ensemble. MBP = grey, actin = black.

AlphaFold3 models were further matched to the SAXS curves (Schneidman-Duhovny et al., 2013). The single best conformer generated by BilboMD with the clustered residues mentioned above bound to actin as a rigid domain (Pelikan et al., 2009) did not match the data (**χ**^2^=9 for MBP-Sca2/6 and **χ**^2^=4 for MBP-Sca2/6 plus actin), indicating the conformational variability of the protein in the solution state. The 2-state fit for both MBP-Sca2/6 and MBP-Sca2/6 plus actin (Figure 4C and D) showed an excellent fit to the data (Figure 4C). MBP-Sca2/6-L101A/K102A was not fitted because a discernible complex with actin was not observed. Consistent with the flexible and extended conformation of both proteins suggested by normalized Kratky plot and P(r) function, the 2-state model predicted a significant contribution from unfolded linkers within MBP-Sca2/6 even in the presence of G-actin (Figure 4D). Together, these data suggest that Sca2/6 lacks larger stably folded domains and exhibits high flexibility, even when bound to G-actin.

Finally, to test the functional importance of the G-actin binding motif, we assessed the actin-nucleation activity MBP-Sca2/6-L101A/K102A using the pyrene actin assembly assay. MBP-Sca2/6-L101A/K102A showed no nucleation activity above background levels at concentrations ranging from 10 - 250 nM, in contrast to the clear nucleation activity of MBP-Sca2/6 over the same concentration range (Figure 3B and C). These data demonstrate that the G-actin binding motif is necessary for the actin nucleation activity of Sca2/6.

### *R. bellii* has a single phase of actin-based motility in mammalian cell lines

Although both *R. bellii* RickA and Sca2/6 are biochemically active, their roles in *R. bellii* actin-based motility have not been determined. Our prior work with *R. parkeri* showed that this species exhibits two phases of motility; an early (<2 hours post infection (hpi)) phase driven by RickA and a late phase (>5 hpi) driven by Sca2 (Reed et al., 2014). To determine if *R. bellii* also exhibits this same biphasic motility pattern, we infected human A549 epithelial or HMEC-1 endothelial cell lines with either *R. parkeri* or *R. bellii* and quantified the frequency at which bacteria assemble actin tails between 15 min post-infection (mpi) and 24 hpi as a measure of the frequency of actin-based motility. Actin tails were observed throughout the 24 h time course in *R. parkeri-*infected cells, with two peaks in actin tail frequency corresponding to the early RickA-motility (15 - 30 mpi) and the late Sca2-motility (16 hpi - 24 hpi) phases (Figure 5A and B). In contrast, *R. bellii* actin tails were extremely rare prior to 8 hpi in both cell lines (Figure 5A and B). The frequency of *R. bellii* actin tails increased from 8 hpi through 24 hpi. These results suggest that *R. bellii* exhibits only a single phase of actin-based motility in infected mammalian cell lines.

**FIGURE 5.**
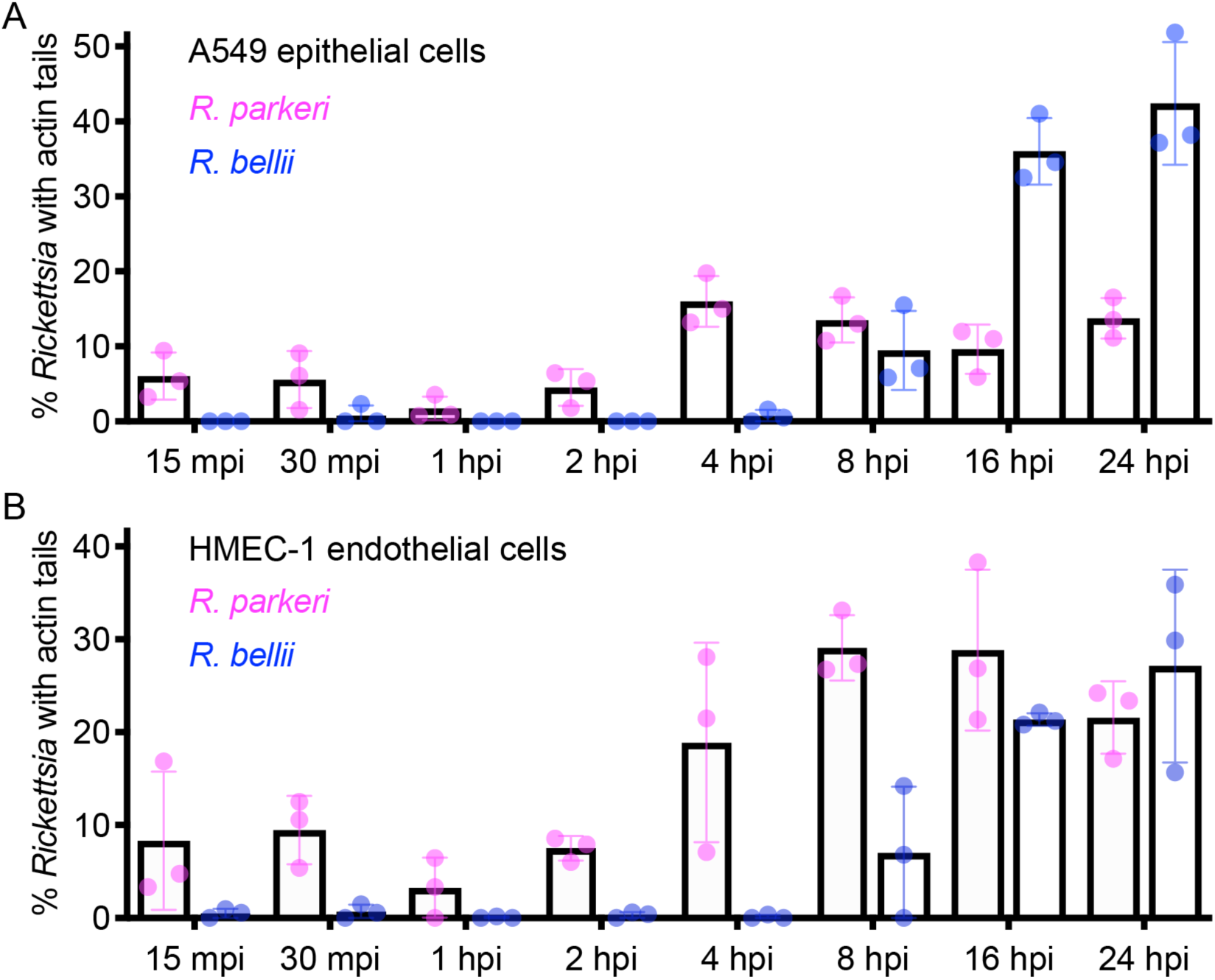
*R. bellii* has a single phase of actin-based motility. Quantification of the percentage of *R. parkeri* (magenta) and *R. bellii* (blue) actin tails in infected (A) A549 human epithelial cells and (B) HMEC-1 human endothelial cells from 15 mpi to 24 h hpi. Error bars indicate SD. N = 3 independent experiments, n ≥ 51 bacteria per experiment per time point.

### No role was observed for *R. bellii* RickA and host Arp2/3 complex in actin-based motility

To investigate a possible role for RickA in *R. bellii* actin-based motility at 8 and 24 hpi, we first localized surface-associated RickA and actin by fluorescence microscopy in infected A549 cells. Although actin tails were observed at both time points, we did not observe RickA on the bacterial surface (Figure 6A and B). Next, to localize intra-bacterial RickA, we permeabilized bacteria with lysozyme. A subset of bacteria showed evidence of internal RickA staining (Figure 6A and B), consistent with our immunoblotting data showing RickA is expressed by *R. bellii* (Figure S3). These data suggest that RickA is not localized to the bacterial surface under the conditions tested and therefore its surface localization is not correlated with actin-based motility.

**FIGURE 6.**
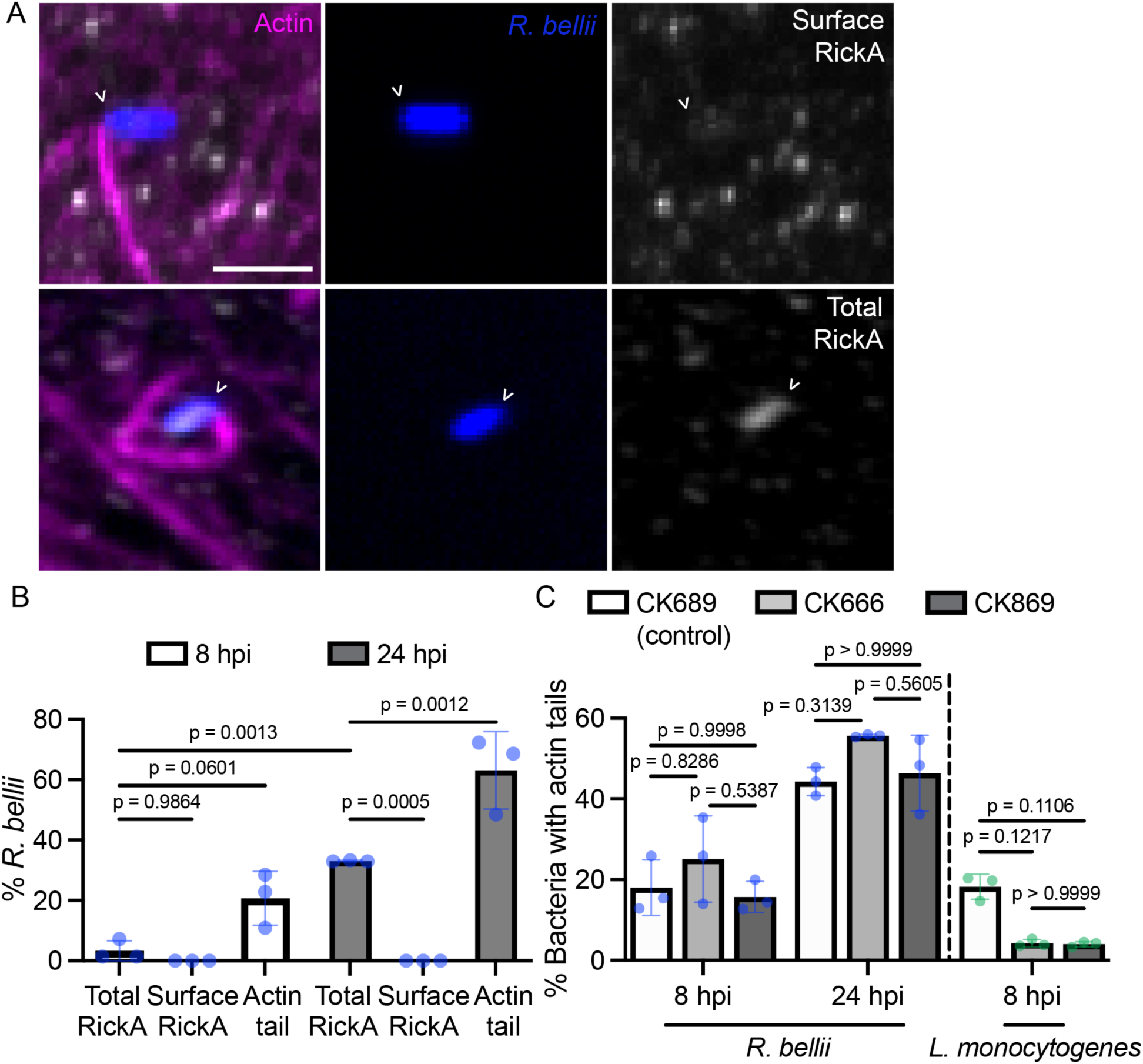
RickA was not observed on the *R. bellii* surface and Arp2/3 complex inhibition did not impact *R. bellii* motility. (A) Representative maximum-projection images of *R. bellii* (blue) in infected A549 cells immuno-stained for surface-localized and total RickA (white) and actin filaments (magenta) at 24 hpi. The bacterial pole associated with an actin tail is indicated with open arrowheads. Scale bar = 1 µm. (B) Percentage of *R. bellii* with total and surface-localized RickA at 8 and 24 hpi. N = 3 independent experiments, n ≥ 95 bacteria per experiment per time point. Significance was determined using a two-way ANOVA with a Tukey’s multiple comparisons post-test. (C) Percentage of *L. monocytogenes* and *R. bellii* with actin tails at 8 and/or 24 hpi in the presence of Arp2/3 complex-inhibitors CK666 or CK869, or inactive control compound CK689. N = 3 independent experiments, n ≥ 139 bacteria per experiment per time point. Error bars indicate SD. Significance was determined using a one-way ANOVA with a Tukey’s multiple comparisons post-test.

To examine the functional importance of RickA, we leveraged the dependency of RickA-motility on the host Arp2/3 complex and the availability of small-molecule Arp2/3 complex inhibitors CK-666 and CK-869, as well as inactive control compound CK-689 (Nolen et al., 2009; Cao et al., 2024). As a positive control, we evaluated the effect of 1 h treatment with these compounds on *L. monocytogenes* actin-based motility in infected A549 cells, a process that is known to depend on the Arp2/3 complex (Welch et al., 1997). As expected, *L. monocytogenes* actin tail frequency was significantly reduced in the presence of CK-666 or CK-869 compared to the inactive control drug (Figure 6C). In contrast, for *R. bellii*-infected A549 cells at either 8 hpi or 24 hpi, actin tail frequencies were not significantly different following 1 h treatment with the Arp2/3 complex inhibitors CK-666 or CK-869 versus the inactive control compound (Figure 6C). Thus, while RickA is biochemically active, RickA- and Arp2/3 complex-dependent actin-based motility was not observed in *R. bellii-*infected human cell lines.

### Sca2/6 surface localization is correlated with *R. bellii* actin-based motility

We next investigated a role for Sca2/6 in *R. bellii* actin-based motility at 8 and 24 hpi by localizing surface-associated Sca2/6 and actin by confocal microscopy in infected A549 cells. Sca2/6 exhibited a focused localization at the poles of ∼40% of bacteria at 8 hpi and ∼80% of bacteria at 24 hpi (Figure 7A and B). A majority of bacteria with polar Sca2/6 foci had an associated actin tail. Moreover, for all bacteria with an actin tail in this data set, the actin tail emerged from the bacterial pole with Sca2/6 (Figure 7A). Therefore, Sca2/6 localization to the surface of *R. bellii* is tightly correlated with actin-based motility.

**FIGURE 7.**
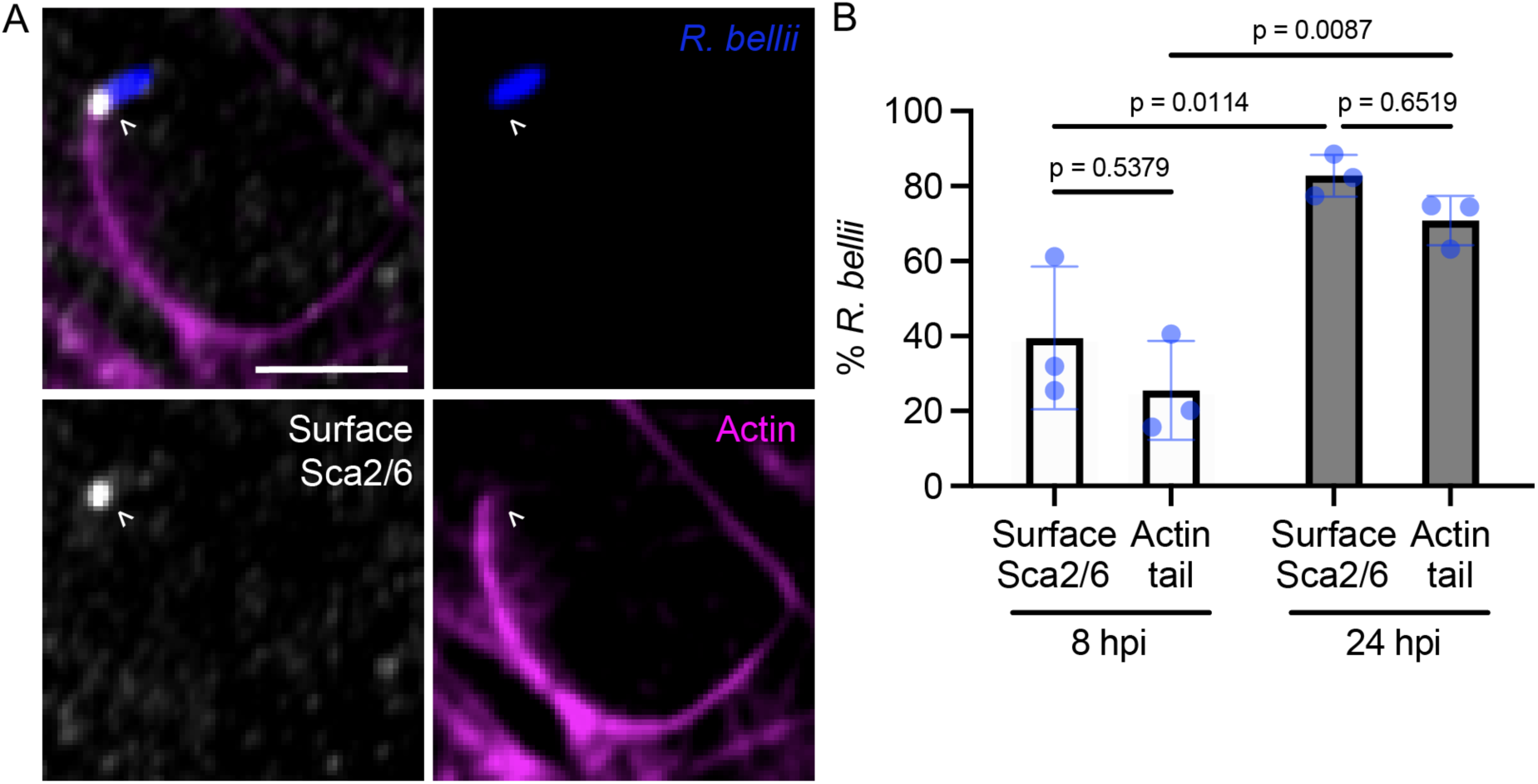
Sca2/6 localizes to the surface of *R. bellii* in correlation with the formation of actin tails. (A) Representative confocal maximum-intensity projection image of *R. bellii* (blue) with actin filaments (magenta) and Sca2/6 (white) in infected A549 cells at 8 hpi. The bacterial pole associated with an actin tail is indicated with open arrowheads. Scale bar = 2 µm. (B) Percentage of *R. bellii* with polar Sca2/6 and frequency of actin tails. N = 3 independent experiments, n ≥ 94 bacteria experiment per time point. Error bars indicate SD. Significance was determined using a two-way ANOVA with a Tukey’s multiple comparisons post-test.

### Sca2 ortholog localization and actin tail architecture differ between *Rickettsia* species

Sca2/6 localization to a discrete focus at the *R. bellii* pole seemed distinct compared with previously reported Sca2 localization in a more cup-shaped pattern at the pole of *R. parkeri* (Haglund et al., 2010; Reed et al., 2014). To directly compare differences in Sca2 ortholog localization patterns on the surface of *R. parkeri* versus *R. bellii*, we imaged *Rickettsia-*infected A549 cells at 24 hpi using a super-resolution lattice structured illumination microscope (LSIM) and further deconvolved the images to reduce image distortion. *R. parkeri* Sca2 tended to coat the bacterial pole associated with an actin tail or even the entire bacterium, while *R. bellii* Sca2/6 localized to the bacterial pole as a distinct punctum (Figure 8A). We further quantified the surface area and sphericity of the Sca2 ortholog signal, along with the overlapped volume between the Sca2 ortholog and the bacterial signals. *R. parkeri* Sca2 occupied a larger surface area (Figure 8B) and exhibited a greater overlap with the bacterium (Figure 8C) compared with *R. bellii* Sca2/6. Sca2/6 localization was significantly more spherical on *R. bellii* than Sca2 localization was on *R. parkeri* (Figure 8D). Therefore, the Sca2 ortholog exhibits a more extreme polar localization on *R. bellii* compared to *R. parkeri*.

**FIGURE 8.**
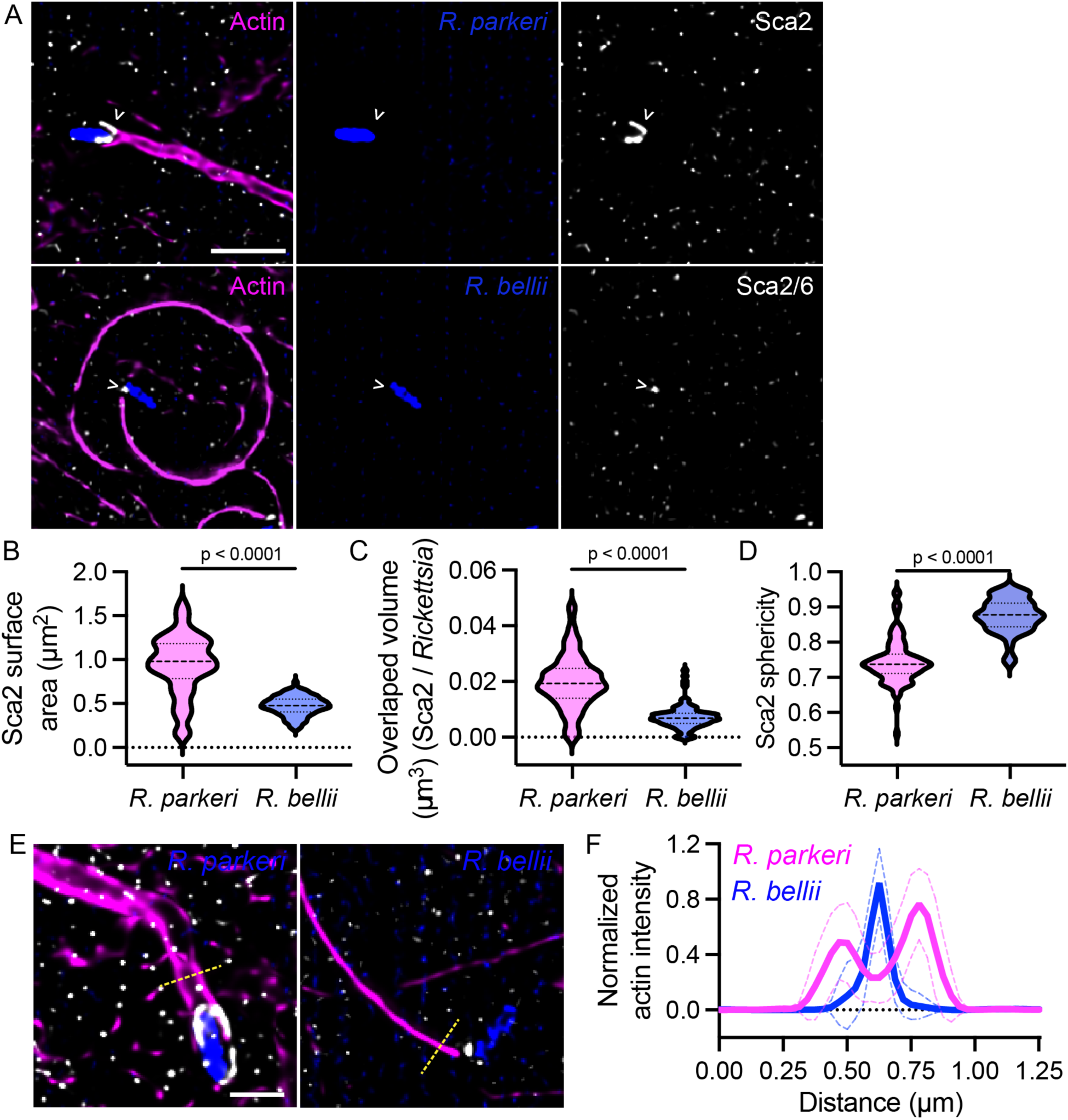
Sca2 ortholog morphology and actin tail organization differs between *R. bellii* and *R. parkeri*. (C) Deconvolved lattice structured illumination (LSIM) maximum projections of A549s infected for 24 h with *R. parkeri* (blue) or *R. bellii* (blue) stained actin filaments (magenta) and Sca2 ortholog (white). Scale bars = 2 µm. Quantification of *R. parkeri* Sca2 and *R. bellii* Sca2/6 (D) surface area and (E) sphericity, and (F) overlapped volume. Error bars indicate SD. N = 3 independent replicates, n ≥ 229 bacteria per species per replicate. Significance was determined using a Welch’s t test. (E) Maximum projections of deconvolved LSIM micrographs showing representative actin tail (magenta) intensity traces (dashed yellow line) positioned 0.5 µm from the start of the actin tail at the pole of the bacterium (blue) with Sca2 ortholog fluorescence (white) in infected A549 cells. Scale bar = 1 µm. (F) Plot of cumulative *R. parkeri* and *R. bellii* actin tail intensity traces. Dashed lines indicate SD. N = 3 independent experiments, n = 18 traces per *Rickettsia* species.

Corresponding with differences in Sca2 ortholog localization, the architecture of the actin tails in the LSIM images was distinct for *R. parkeri* and *R. bellii* at 24 hpi, when actin-based motility is predominant for both species. *R. parkeri* actin tails typically had two bands of actin emanating from the sides of the bacterial pole (Figure 8A and E), whereas *R. bellii* actin tails had a single band of actin centered on the pole (Figure 8A and E). To quantify the spatial organization of actin tails, we took a line intensity scan of actin fluorescence perpendicular to the long axis of the actin tail (0.5 µm from the bacterial pole). *R. parkeri* actin tails showed two distinct intensity peaks spaced 0.30 ± 0.09 µm apart, whereas *R. bellii* actin tails showed a single intensity peak (Figure 8F). The correlation between Sca2 ortholog localization and actin intensity profiles suggest that differences in protein localization may impact differences actin tail organization between *Rickettsia* species.

### Differences in Sca2 ortholog mechanism and localization, as well as host cell type, correlate with differences in actin-based motility

We lastly looked at the functional consequences of differences in Sca2 ortholog mechanism and localization on actin-based motility parameters. We carried out live-cell imaging of *R. parkeri*- or *R. bellii*-infected A549 or HMEC-1 cells stably expressing F-tractin-TagRFPT (Johnson and Schell, 2009) at 24 hpi for 5 min and analyzed the actin-based motility speed and efficiency for each *Rickettsia* species and host cell combination. *R. parkeri* motility was significantly faster in A549 cells (mean 21 ± 5 µm/min) compared with HMEC-1 cells (mean 11 ± 4 µm/min). *R. bellii* moved more slowly than *R. parkeri* and with indistinguishable rates in A549 (mean 8 ± 3 µm/min) versus HMEC-1 cells (mean 9 ± 3 µm/min) (Figure 9A, Movie S2). We also measured the motility efficiency (defined as the straight-line distance between the starting and stopping points (Δd) in a 1 min interval, divided by the total distance traveled (D) during the interval, or Δd/D). *R. parkeri* motility efficiency was higher in both host cell lines (A549 mean 0.8 ± 0.2; HMEC-1 mean 0.8 ± 0.2) than for *R. bellii* (A549 mean 0.6 ± 0.3, HMEC-1 mean 0.7 ± 0.2) (Figure 9B, Movie S3). Traces of representative bacterial motility paths revealed more curved paths for *R. bellii* than for *R. parkeri*, particularly in A549 cells (Figure 9C). Thus, differences in Sca2/6 mechanism and localization correlate with slower *R. bellii* speed and lower motility efficiency compared with *R. parkeri*.

**FIGURE 9.**
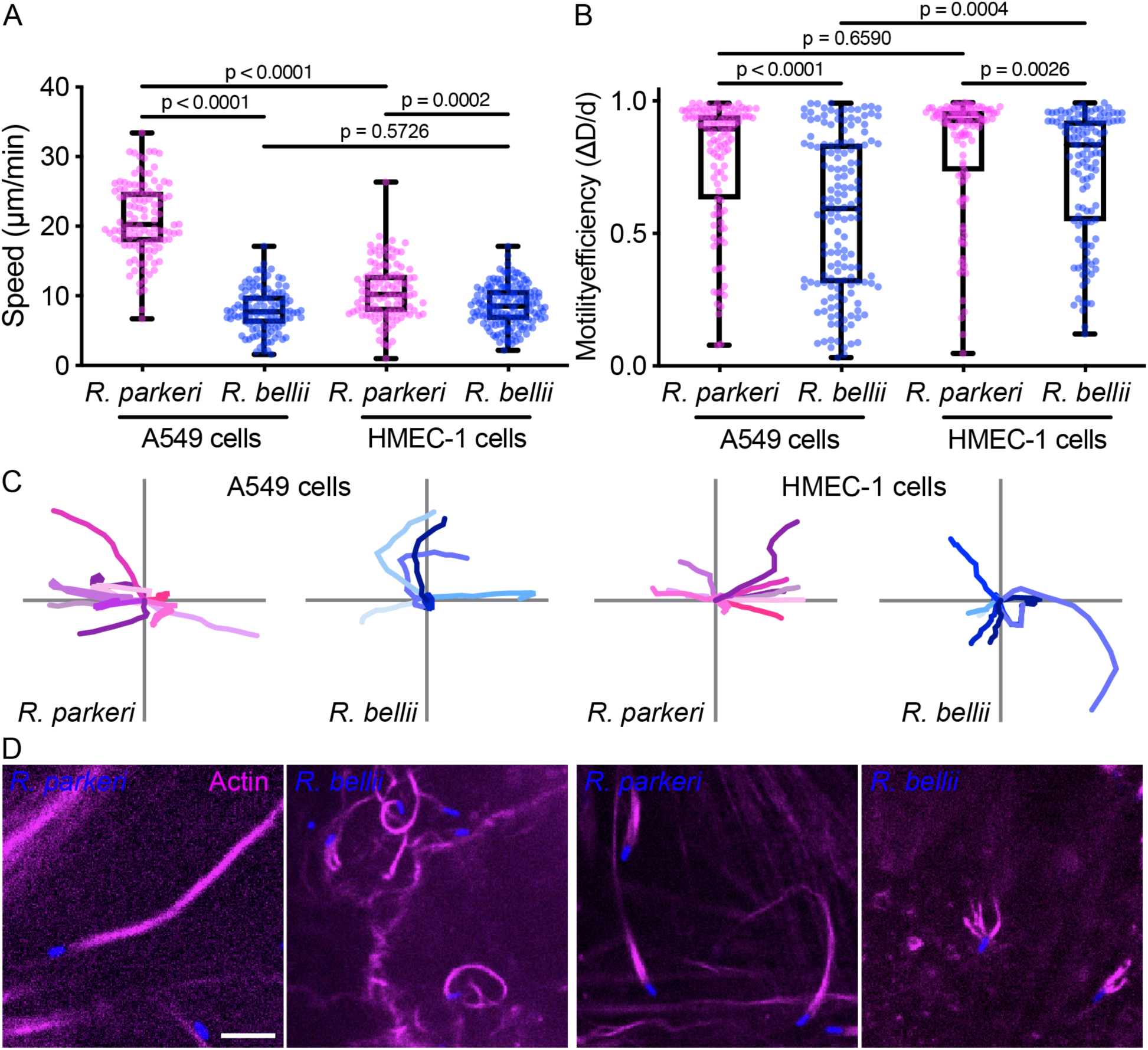
Actin-based motility parameters and actin tail organization differ between *R. bellii* and *R. parkeri* as well as host cell lines. (A) Speed of *R. parkeri* and *R. bellii* at 24 hpi in A549 and HMEC-1 cell lines. (B) Motility efficiency (ΔD/d) of *R. parkeri* and *R. bellii* at 24 hpi in A549 and HMEC-1 in 1 min intervals. N = 3 - 5 independent experiments per *Rickettsia* species and host cell line, n ≥ 110 bacteria per *Rickettsia* species and host cell line. Significance was determined using a one-way ANOVA with a (A) Tukey’s multiple comparisons or (B) Kruskal-Wallis post-test. (C) Motility traces of 10 representative *R. parkeri* (magenta) and *R. bellii* (blue) in infected A549 or HMEC-1 cell lines. Some traces are not clearly visible due to overlapping tracks. (D) Representative still images of *R. parkeri* and *R. bellii* (blue) with actin tails (magenta), showing differences in actin organization. Scale bar = 5 µm.

Notably, the organization of *R. bellii* actin tails was dramatically different in A549 versus HMEC-1 cells. In A549 cells, actin tails were consolidated, consisting of a single bundle of actin filaments (Figure 8A and 8D). In contrast, in HMEC-1 cells, *R. bellii* actin tails were disorganized, consisting of multiple bundles of actin filaments splaying from the bacterial pole (Figure 9D). As noted above, *R. parkeri* actin tails appeared similar in organization in the different cell types (Figure 8A and 9D). Thus, differences in Sca2 mechanism and localization also correlate with cell type-specific differences in *R. bellii* actin-tail organization.

## Discussion

Our results provide insight into the evolution of actin-based motility effectors within the *Rickettsia* genus. We find that RickA orthologs have maintained the ability to activate the host Arp2/3 complex to nucleate actin filaments, but RickA function in actin-based motility is not conserved between species. Sca2 paralogs, on the other hand, contribute to actin-based motility in divergent *Rickettsia* species, but their sequences and mechanisms differ substantially between species. Divergence of nucleation mechanisms between Sca2 orthologs coupled with different localization patterns on the bacterial surface contribute to differences in actin organization and motility. Our data indicate there is considerable evolutionary plasticity in actin-based motility regulation and mechanisms within the *Rickettsia* genus.

In *R. parkeri*, RickA-motility was previously shown to occur early after infection of a host cell and to function in cell-cell spread (Reed et al., 2014; Tran et al., 2025). Although we now report that *R. bellii* bacteria express a RickA ortholog that possesses Arp2/3-dependent actin nucleation activity, we found no evidence for RickA localization to the surface of *R. bellii*, or RickA and Arp2/3 complex function in *R. bellii* actin-based motility. The conservation of RickA in divergent species (Lehman et al., 2024), despite the genus undergoing reductive genome evolution (Blanc et al., 2007), suggests that RickA performs an important function. One explanation for our observations is that the function and/or regulation of RickA is host dependent. SFG I species, such as *R. parkeri*, have adapted to use RickA-motility in tick cell lines, where an *R. parkeri rickA* transposon mutant showed reduced cell-cell spread (Harris et al., 2018), as well as in mammalian cell lines and animals, where the same *rickA* mutant was deficient in cell-cell spread and eschar formation (Tran et al., 2025). In contrast, the tick endosymbiont *R. bellii* may only use RickA in tick cells. Our observation that RickA is expressed in both *R. parkeri* and *R. bellii* but is only secreted and localized to the surface of *R. parkeri*, suggests potential differences in the regulation of RickA secretion. It also remains possible that RickA performs important functions outside of actin-based motility.

On the other hand, Sca2 orthologs are quite different between divergent *Rickettsia* species. The *R. bellii* genome contains a single *sca2/6* gene, whereas in other species this gene appears to have undergone a tandem duplication event, yielding adjacent *sca2* and *sca6* genes that are intact in some species and degraded in others (Blanc et al., 2005; Ngwamidiba et al., 2005; Sears et al., 2012). Gene duplication can contribute to more rapid evolution (Zhang, 2003) and may have contributed to the evolution of the passenger domains of Sca2 orthologs, which differ considerably in primary sequence. Previous studies showed that Sca2 from SFG I species nucleates and caps actin in the absence of profilin, enables elongation, and processively associates with growing barbed ends in the presence of profilin, similar to eukaryotic formins (Haglund et al., 2010; Madasu et al., 2013; Alqassim et al., 2019; Carman et al., 2023). SFG I Sca2 also assumes a folded formin-like structure, with the N-terminal region forming ring-shaped structural core in conjunction with the CRD (Alqassim et al., 2019; Carman et al., 2023). In contrast, we report that *R. bellii* Sca2/6 nucleates and modestly enhances elongation in the absence of profilin and does not cap or stably associate with filament barbed ends, characteristics that differ from eukaryotic formins. Moreover, Sca2/6 is structurally dissimilar from SFG I Sca2 and eukaryotic formins, as SAXS analysis indicates that it lacks larger stably folded domains and exhibits high flexibility. SAXS analysis coupled with AlphaFold 3 predictions also indicate that *R. bellii* Sca2/6 complexes with G-actin primarily via amino acids 70-110 and exhibits weaker interactions with G-actin via amino acids 110-180. Thus, motifs interacting with G-actin comprise much of the N-terminal region of the protein. Although the G-actin binding domain between amino acids 70-110 is necessary for actin nucleation, the broader region of the protein between amino acids 70-180 is not sufficient, and the CTD is also required. The contribution of the CTD to biochemical activity may be through potential low affinity interactions with F-actin. Only the poly-proline motif is dispensable for nucleation activity *in vitro*, although poly-proline sequences bind profilin (Tanaka and Shibata, 1985) and thus may contribute to actin assembly in host cells. Moreover, while SAXS analysis reveals that Sca2/6 is a monomer in solution, crowding on the bacterial surface may also affect actin nucleation and elongation activity.

The sequence and biochemical properties of *R. belliii* Sca2/6 differ from other microbial actin nucleators. Sca2/6 possesses a single higher affinity G-actin-binding domain predicted to bind the barbed end of an actin monomer and does not exhibit detectable association with either end of actin filaments. These properties differ from the tandem WH2-domain nucleation factors VopL/VopF from *Vibrio* species, which have three WH2 domains spaced by linkers and have been shown to dimerize for efficient actin nucleation and elongation via the pointed or barbed end of an actin filament (Tam et al., 2007; Liverman et al., 2007; Yu et al., 2011; Namgoong et al., 2011; Zahm et al., 2013; Avvaru et al., 2015; Burke et al., 2017; Kudryashova et al., 2022). *R. bellii* Sca2/6 also lacks key characteristics of the host ENA/VASP-like protein BimA from *Burkholderia* species (Benanti et al., 2015), such as the ability to trimerize. Based on our current understanding, *R. bellii* Sca2/6 therefore exhibits a distinctive mechanism of actin nucleation.

Our comparison of Sca2 ortholog localization in *R. parkeri* and *R. bellii* also reveals clear differences, with Sca2/6 in *R. bellii* exhibiting a more extreme polar distribution than Sca2 in *R. parkeri*. Differences in localization may be a result of difference in protein secretion or processing. The *Shigella* actin-based motility effector IcsA is also a T5SS protein that is secreted at the bacterial pole and can diffuse across the surface, where it is cleaved by a protease (Shere *et al*., 1997; Steinhauer *et al*., 1999). When polar secretion is rapid, slower cleavage across the entire bacterial surface acts as a pruning mechanism to keep IcsA restricted to the bacterial pole (Shere et al., 1997; Steinhauer et al., 1999). *R. parkeri* Sca2 is also cleaved (Haglund et al., 2010; Reed et al., 2014), as is *R. bellii* Sca2/6 (this study). Variations in location/rates of autotransporter secretion and protease activity may explain why the Sca2 ortholog covers more of the bacterial surface of *R. parkeri* compared with *R. bellii*.

At time points when Sca2 is the predominant motility effector for both *R. parkeri* and *R. bellii*, we observed clear differences in actin filament organization and actin-based motility. *R. parkeri* actin tails are organized into two discrete bundles that emerge from the sides of the bacterial pole, similar to prior reports (Van Kirk et al., 2000; Haglund et al., 2010), and bacteria move more quickly and in straighter trajectories. *R. bellii* actin tails are organized into a single bundle, and bacteria move more slowly and in more meandering trajectories. Variations in motility likely derive from differences in Sca2 ortholog mechanisms of action and localization patterns. Differences in biochemical activity may affect rates of actin elongation and/or filament capping at the bacterial surface. The fuller coverage of *R. parkeri* Sca2 on the bacterial surface and the resulting actin density that brackets the bacterial pole may steer the bacteria in straighter trajectories. In contrast, the extreme polar distribution of *R. bellii* Sca2/6 and the reduced actin density may contribute to more meandering motility and a smaller number of elongating actin filaments generating propulsive force. We also observed host cell type-specific differences, with disorganized *R. bellii* actin tails consisting of multiple splayed filament bundles in human endothelial cells. This actin tail architecture is reminiscent of the splayed *R. parkeri* actin tails observed in infected cells depleted of the actin-bundling protein fimbrin by siRNA (Serio et al., 2010). *R. parkeri* bacteria, which naturally target human endothelial cells during infection (Scott et al., 2022), retained a more consolidated actin tail phenotype in the endothelial cell line. Thus, differences in Sca2 ortholog localization, along with differences in the abundance and recruitment of host actin-bundling proteins or other actin-associated proteins, likely play an important role in defining actin tail organization and bacterial movement parameters.

Together, our results provide insight into the evolution of actin-based motility mechanisms in microbial pathogens. While divergent species throughout the *Rickettsia* genus maintain the ability to undergo actin-based motility, the utilization of orthologs of the two motility effectors RickA and Sca2, and the biochemical mechanisms of Sca2 ortholog-mediated actin polymerization, have diverged. This evolution likely reflects changes in both motility effector sequences and regulation. Because of the diversity of *Rickettsia* genus, these bacteria provide a unique platform to unravel how actin-based motility evolved and contributes to pathogenicity or colonization. Future studies of how actin-based motility is affected by motility effector expression and localization, actin ultrastructure and organization, and the host cell environment will provide powerful insights into both pathogen infection and host cell biology.

## Materials and methods

### Mammalian cell culture and cell line generation

Vero (CCL-81), HEK293T (CRL-3216), A549 (CRM-CCL-185), and HMEC-1 (CRL-3243) cells were sourced from the University of California, Berkeley, tissue culture facility. Vero cells were grown in DMEM containing 4.5 g/L D-glucose, 4 mM L-glutamine (Gibco, 11965118) and 2% fetal bovine serum (Gemini Bio-products, 100-500). HEK293T and A549 cells were grown in DMEM containing 4.5 g/L D-glucose, 4 mM L-glutamine (Gibco, 11965118), and 10% fetal bovine serum (ATLAS Biologics, FP-0500-A). HMEC-1 cells were grown in MCDB 131 (Sigma, M8537) containing 10 mM L-glutamine (Sigma, M8537-1L), 14 mM sodium bicarbonate, 10% fetal bovine serum (Hyclone, SH30071.03), 10 ng/ml EGF (BD Biosciences, 3540001), and 1 µg/ml hydrocortisone (Sigma, H0888). All cells were grown at 37°C in 5% CO_2_.

A549 or HMEC-1 cells expressing F-tractin-TagRFPT were generated through retroviral transduction with SC1205 as previously described (Lamason et al., 2016). SC1205 (F-tractin-TagRFPT-pIPFCW2) was generated by subcloning the genes encoding F-tractin (flanked by NheI and XhoI restriction sites) (Johnson and Schell, 2009) and TagRFPT (flanked by XhoI and EcoRI restriction sites) (Shaner et al., 2008) into NheI- and EcoRI-digested pIRES-puro (IP)-FCW2. Transduced cells were selected with 1 µg/mL puromycin (Sigma, P8833) and sorted using the Aria Fusion Cell Sorter in the UC Berkeley Flow Cytometry Facility to select for cells that fell in the top 30% of brightness for RFP fluorescence.

### Bacterial propagation and strain construction

*R. parkeri* strain Portsmouth was obtained from Chris Paddock at the Centers for Disease Control and prevention. *R. bellii* strain RML 369-C was obtained from Ted Hackstadt at the NIH/NIAID Rocky Mountain Laboratories. For propagation, *Rickettsia* bacteria were allowed to infect 5 confluent T175 flasks of Vero cells for 4-7 d at 33°C until cells rounded and started to dissociate from the monolayer. Infected cells were scraped from the bottom of the flask into the growth media, pelleted, resuspended in 10 ml of K36 buffer (50 mM KH_2_PO_4_, 50 mM K_2_HPO_4_, pH 7.0, 100 mM KCl, 15 mM NaCl). Bacteria were released on ice from Vero cells by Dounce homogenization (50 strokes). Bacteria were isolated by centrifugation in a solution of 30% RenoCal-76 (Bracco Diagnostics, 04H208) in K36 buffer using a SW-32 Ti rotor (Beckman/Coulter) at 18,500 rpm for 30 min at 4°C. The bacterial pellet was resuspended in 2 mL brain heart infusion (BHI) media (BD Difco, DF0418-17-7) and aliquoted before freezing at - 80°C.

Bacterial titers were determined by serially diluting the bacterial stock and performing a plaque assay in Vero cells. Briefly, serial dilutions in Vero cell media were used to infect a 6-well plate of Vero cells by centrifugation at 300 x g for 5 min at room temperature and overnight incubation at 33°C. Cells were then overlaid in DMEM with 10% FBS (Gemini Bio-products, 100-500) and 0.5% (w/v) agarose (Sigma A9539). Once plaques appeared (5-7 d post-infection) cells were overlaid with a mixture of 1x PBS (Gibco, 10010072), 0.5% agarose, and 2.5% (v/v) neutral red (Sigma, N6264). Plaques were quantified the next day and used to calculate the plaque forming units (PFU) per mL.

Fluorescent *R. parkeri* or *R. bellii* expressing pRAM18dRGA (Burkhardt et al., 2011) or pRAM18dR-2xCoTagBFP (Figueroa-Cuilan et al., 2023) were generated by infecting a T75 flask of Vero cells with bacteria and incubating at 33°C. Infected Vero cells were scraped up, resuspended in K36 buffer, and then broken open with 1 mm beads by two 30 s rounds of vortexing and resting on ice. Cell debris was pelleted, and the supernatant was washed 3x with 250 mM sucrose (ThermoFisher, S5). Washed bacteria were resuspended in 100 µL BHI and incubated on ice for 20 min with 10 µg of the plasmid. Bacteria were then electroporated in a 0.2 cm cuvette at 2.5 kV, 200 Ω, 25 µF. Electroporated bacteria were resuspended in 1.2 mL BHI and aliquoted onto Vero cells in 6-well plate (200 uL per well). The plate was centrifuged at 300 x g for 5 min at room temperature, and then incubated at 33°C. The next day, 500 ng/mL rifampicin (Sigma, R3501; resuspended to 1 mg/mL in methanol and filter sterilized) was added to each well of infected Vero cells. After clear signs of infection were observed, *Rickettsia* were harvested by adding cold sterile water to each well for 2 min to break bacteria out of Vero cells. An equal amount of 2x BHI was added to each well and bacteria were frozen at −80°C. After visually confirming fluorescent protein expression, bacteria were propagated as described above.

*L. monocytogenes* strain 10403S::dasherGFP was obtained from Daniel A. Portnoy at the University of California, Berkeley. *L monocytogenes* was streaked from frozen stocks onto BHI agar plates with 200 µg/mL spectinomycin and grown overnight at 37°C. For infections, single colonies were used to seed overnight cultures in BHI with 200 µg/mL spectinomycin and were grown at 30°C at a slant with no shaking.

### Generating plasmids for protein expression

To generate plasmids pET-GST-RickA and pET-MBP-RickA to express *R. bellii* RickA tagged with glutathione-S transferase (GST) or maltose binding protein (MBP), the *R. bellii rickA* gene was amplified from boiled bacteria by PCR (primers: 5’-CTGTACTTCCAATCCAATGCAAAGATAACTGAGCTT-3’ and 5’-CCGTTATCCACTTCCAATCTACCTTTGTTGAGATTGTT-3’). To generate the plasmid pET-MBP-Sca2/6 to express *R. bellii* Sca2/6 tagged with MBP, the sequence encoding the Sca2/6 passenger domain (aa 43-630) excluding the secretion signal sequence and autotransporter domain was amplified from plasmid DNA by touchdown PCR (Korbie and Mattick, 2008) (primers: 5’-ACCTGTACTTCCAATCCAATGCACCACCTCCACCACCC_3’ and 5’-ATCCGTTATCCACTTCCAATTCATTCATCACCACCAGTAATCGCAG-3’). The primers used to amplify the *rickA* and *sca2/6* genes included homology arms that facilitated Gibson cloning (New England Biolabs, E2611S) into SspI-cut plasmids pET His6 MBP TEV LIC (Addgene #29656) or pET His6 GST TEV LIC (Addgene #29655) obtained from University of California, Berkeley, QB3 MacroLab.

To generate plasmids pET*-*MBP-Sca2/6-NTD, pET-MBP-Sca2/6-ΔpolyP, and pET-MBP-Sca2/6-CTD to express Sca2/6 truncation derivatives, desired regions of the plasmid pET-MBP-Sca2/6 were amplified by PCR (primers: pET-MBP-Sca2/6-NTD: 5’-TGGATTAAAATTAGAAATATAAGATG-3’ and 5’-TTTGCACTTGCTTCCCCCAC-3’; pET-MBP-Sca2/6-ΔpolyP, 5’-GTTGATCCTAAATCAGCAGC-3’ and 5’-CATGAACTATATCTCCTTCTTAAAG-3’; pET-MBP-Sca2/6-CTD, 5’-AATTATAAGTATATAATTTCGTATAGTACG-3’ and 5’-CATGAACTATATCTCCTTCTTAAAG-3’). To generate plasmids pET-MBP-Sca2/6-L101A/K102A, pET-MBP-Sca2/6 was amplified with primers encoding the L101A/K102A mutations (primers: 5’-TTATGATACAGCGGCGAAAAATAGGAAAAATATAAAAAATAAAAAG-3’ and 5’-GGATTAAATGTTGGACCTAATTC-3’). To generate pET-MBP-Sca2/6-KCK, pET-MBP-Sca2/6 was amplified by PCR with primers encoding a C-terminal GGS linker and KCK motif (primers: 5’-AGCAAGTGCAAATGAATTGGAAGTGGATAAC-3’ and 5’-TCCCCCACCGCTTCCACCGCCTTCATCACCACCAGTAATC-3’). The resulting PCR products where ligated using the KLD reaction mix (New England Biolabs, M0554S).

To generate plasmids pGEX-GST-Sca2/6-44 (aa 44-175) and pGEX-GST-Sca2/6-357 (and aa 357-464) for anti-Sca2/6 antibody production, gene sequences encoding the *R. bellii* Sca2/6 truncations were subcloned into pGEX-6P-3 (GE Healthcare, 28-9546-51).

### Protein purification and fluorescent labeling

To express *R. bellii* GST-RickA and MBP-RickA, BL21-CodonPlus (DE3)-RIL *E. coli* (Agilent Technologies, 230245) were transformed with plasmid pET-GST-RickA or pET-MBP-RickA. For protein expression, bacteria were grown overnight at 37°C in LB media. The overnight culture was then diluted 1:100 into 3 L of 2xYT media and grown to an OD_600_ of 0.5-0.8. Protein expression was induced by adding IPTG to 0.3 mM and bacteria were grown at 37°C for an additional 2 h. Cells were pelleted at 4000 rpm for 20 min at 4°C, resuspended in 30 mL either GST lysis buffer (50 mM Tris, pH 7.5, 250 mM KCl, 1 mM EDTA) or MBP lysis buffer (20 mM HEPES, pH 8, 200 mM NaCl, 1 mM EDTA) with 1 mg/mL lysozyme (Sigma, L4919) and protease inhibitors (10 µg/mL leupeptin, pepstatin, chymostatin (LPC) and 1 mM PMSF), and lysed by sonication (8-25 x 10s pulses, 30% amplitude). The lysate was centrifuged at 13,000 rpm in a Sorvall SS-34 rotor for 30 min at 4°C. Lysate supernatant was run over Glutathione Sepharose 4B resin (GE Healthcare) or Amylose resin (New England Biolabs), washed with MBP or GST column buffer (the respective lysis buffer with 0.5 mM DTT), and eluted in GST column buffer with 10 mM glutathione or MBP column buffer with 10 mM maltose. MBP-RickA was used for antibody affinity purification as described below. GST-RickA was further purified by gel filtration chromatography on a Superdex 200 Increase 10/300 GL column (Cytiva) equilibrated with Buffer A (20 mM HEPES, pH 7.5, 200 mM KCl, 2 mM EGTA, 2 mM EDTA, 0.5 mM DTT, 10% glycerol), then flash frozen in liquid N_2_ and stored at −80°C.

To express GST-Sca2/6-44 and GST-Sca2/6-357 for *R*. *bellii* Sca2/6 antibody generation, BL21-CodonPlus(DE3)-RIPL (Agilent Technologies, 230280) transformed with pGEX-GST-Sca2/6-44 or pGEX-GST-Sca2/6. Bacteria were grown overnight at 37°C in 2xYT media, diluted 1:100 into 3 L of 2xYT media, and then grown again to an OD600 of 0.5-0.8. Protein expression was induced with 0.3-1 mM IPTG (Gold Biotechnology, 20-109) at 16°C overnight. Cells were pelleted, resuspended in lysis buffer (50 mM Tris, 200 mM KCl, 1 mM EDTA, pH 7.5,) with 1 mg/mL lysozyme (Sigma) and protease inhibitors (10 µg/mL LPC and 1 mM PMSF), and lysed by sonication. Lysate was run over Glutathione Sepharose 4B (GE Healthcare) and washed with wash buffer (lysis buffer plus 0.5 mM DTT). PreScision Protease was added to the wash buffer for 16-18 h at 4°C to cleave the GST tag, and cleaved protein was eluted with 1 column volume of wash buffer. Elutions were run over a Superdex 75 10/300 GL column (GE Healthcare) in 20 mM Tris, pH 7.5, 100 mM KCl, 1 mM EGTA, 2 mM EDTA, 0.5 mM DTT, 10% glycerol, then flash frozen and stored at −80°C.

To express the *R. bellii* MPB-Sca2/6 proteins for use in actin assembly assays, BL21-CodonPlus (DE3)-RIPL *E. coli* cells (Agilent Technologies, 230280) transformed with expression plasmids pET-MBP-Sca2/6, pET-MBP-Sca2/6-ΔpolyP, pET-MBP-Sca2/6-L101A/K102A, pET-MBP-Sca2/6-NTD, or pET-MBP-Sca2/6-CTD. Growth and expression conditions were as described above for expression at 16°C. Cells were pelleted, lysed with lysis buffer (50 mM HEPES, pH 7.5, 500 mM KCl, 10 mM imidazole, 10 % glycerol) with 1 mg/mL lysozyme (Sigma) and protease inhibitors (GoldBio ProBlock Gold Bacterial Protease Inhibitor Cocktail, GB-108-2, Roche cOmplete EDTA-free Protease Inhibitor Cocktail, 4693159001, or APEX Protease Inhbitor Cocktail EDTA-free, K1007) followed by sonication (8-25 x 10s pulses, 30% amplitude). Lysate was run over an Ni-NTA resin (Qiagen), washed with column buffer (50 mM HEPES, pH 7.5, 500 mM KCl, 20 mM imidazole, 1 mM DTT, 10% glycerol), and eluted in elution buffer (50 mM HEPES, pH 7.5, 500 mM KCl, 250 mM imidazole, 1mM DTT, 10% glycerol). Elutions were run over a HiTrapQ HP column (Cytiva, 17-1153-01) with a 100 – 800 mM KCl gradient in anion exchange buffer (20 mM Tris, pH 7.5, 10% glycerol, 0.5 mM DTT) and then a Sephacryl S300 or Superdex 200 Increase 10/300 GL column (Cytiva) equilibrated with Buffer B (20 mM HEPES, pH 7.5, 500 mM KCl, 2 mM EGTA, 2 mM EDTA, 0.5 mM DTT, 10% glycerol). Anion exchange was not used when prepping the MBP-Sca2/6-NTD protein. Proteins were flash frozen in liquid N_2_ and stored at −80°C.

To fluorescently label *R. bellii* Sca2/6 (Sca2/6-AF647), MBP-Sca2/6-KCK was expressed as described for MBP-Sca2/6 proteins and purified as described for MBP-RickA. Post purification, the MBP-tag was cleaved with His-TEV protease for 1 h at room temperature and separated on a Superdex 200 Increase 10/300 GL column (Cytiva) equilibrated with Buffer A. Sca2/6-KCK was dialyzed overnight into Buffer A with 1 mM TCEP (Sigma, 646547) instead of DTT. Equimolar amounts of Sca2/6-KCK (35 µg) and AlexaFluor 647-maleimide (Invitrogen, A23047) were mixed and the maleimide reaction was allowed to proceed overnight at 4°C. Excess dye was removed by dialysis into Buffer A before Sca2/6-AF647 was flash frozen in liquid N_2_ and stored at −80°C.

### Antibody production, purification, and immunoblotting

To generate antiserum to *R. bellii* Sca2/6 (anti-RbSca2/6), Sca2/6-44-175 and Sca2/6-357-464 were expressed and purified as described above, mixed in equimolar amounts, and used to immunize two rabbits (Pocono Rabbit Farm & Laboratory, Inc). The generation of antiserum to *R. parkeri* Sca2 (anti-RpSca2; (Haglund et al., 2010)) and antiserum to *R. rickettsii* RickA (anti-RrRickA; (Jeng et al., 2004)) were described previously. To affinity purify anti-RbSca2/6 or anti-RickA antibodies, 2.5 mg of *R. bellii* Sca2/6 truncations (equimolar amounts) or 500 µg of *R. bellii* MBP-RickA were conjugated to 1 ml of NHS-activated Sepharose 4 Fast Flow (GE Healthcare) in ligand-coupling buffer (200 mM NaHCO_3_, pH 8.3, 500 mM NaCl) for 30 min at room temperature or overnight at 4°C. Ligand-coupled resin was incubated with 10 ml of anti-RbSca2/6 serum or anti-RrRickA serum (*R. rickettsii* RickA is similar in sequence to *R. bellii* RickA). Serum was diluted in 10 ml binding buffer (20 mM Tris, pH 7.5) for 1 h at room temperature or overnight at 4°C. Antibodies were eluted using 100 mM glycine, pH 2.5, and fractions were pooled and dialyzed overnight at 4°C against PBS (137 mM NaCl, 2.7 mM KCl, 10 mM Na_2_HPO_4_, 1.8 mM KH_2_PO_4_, pH 7.4) for anti-RickA or TBS (150 mM NaCl, 50 mM Tris-Cl, pH 7.6) for anti-RbSca2/6. Antibodies were concentrated to 0.72 mg/mL (anti-RbRickA) and 2 mg/mL (anti-RbSca2/6) before flash freezing and storage at −80°C, or mixed with 50% glycerol and stored at −20°C.

For immunoblotting, 5×10^6^ of WT *R. bellii* from a 30% prep was diluted 1:1 with 2x Laemmli buffer (Bio-Rad, 1610737) and boiled for 10 min. The samples were run on a 10% SDS-PAGE gel and transferred to a PVDF membrane (Millipore, IPFL00010). The membrane was blocked in TBS-T (20 mM Tris, pH 8, 150 mM NaCl, 0.1% Tween 20) with 5% non-fat dry milk (Apex, 20-241) for 1 h. The membrane was then incubated with 1:1000 anti-RbSca2/6 or anti-RbRickA in TBS-T with 5% milk, washed 3x in TBS-T, and incubated for 1 h with 1:5000 mouse anti-rabbit HRP secondary antibody (Santa Cruz Biotechnology, sc-2357) in TBS-T with 5% milk, and washed again 3x with TBS-T. Following developing with the ECL HRP substrate kit (Advasta, K-12045-D20) for 45 s, bands were imaged on a ChemiDoc MB Imaging System (Bio-Rad).

### Bulk actin assembly assays

Rabbit skeletal muscle actin (Cytoskeleton, AKL99), pyrene labeled actin (Cytoskeleton, AP05), and rhodamine actin (Cytoskeleton, AR05) were dialyzed for 2-3 d in G-buffer (5 mM Tris, pH 7.5, 0.2 mM CaCl_2_, 0.2 mM ATP, 0.5 mM DTT). The dialyzed actin was centrifuged at 100,000 x g in a TLA100 rotor (Beckman Coulter) using an Optima TLX Ultracentrifuge (Beckman Coulter) for 2 h at 4°C. The top 3/4 of the supernatant was collected and run over a Superdex 200 Increase 10/300 GL column (Cytiva) equilibrated with G-buffer. The fractions corresponding to the latter half of the actin peak (the smaller Stokes radius) were combined and used. Purified Arp2/3 complex (Cytoskeleton, RP01P) was resuspended in milliQ water according to the manufacturer instructions.

Pyrene actin assembly assays were performed by combining 1 µM G-actin mix (10% pyrene labeled) with purified proteins, Buffer A, 10x initiation buffer (10 mM MgCl_2_, 10 mM EGTA, 5 mM ATP, 500 mM KCl) with a final KCl concentration of 96 mM. Fluorescence was measured in two replicates per reaction condition, in three independent experiments, using a Tecan Infinite F200 Pro plate reader and Magellan software v7.1 at 20 s intervals with a 365 nm excitation and 405 nm emission filters. Each curve was baseline-corrected using an average of the first 3 values and graphed in Prism 10.3.1 (GraphPad Software, Inc).

For maximum slope calculations, the first derivative was calculated after smoothing (4 neighbors on each side, 0^th^ order polynomial) and averaging raw fluorescence values of two biological replicates for each sample condition using Prism 10.3.1. *R. bellii* Sca2/6 maximum slope values were then normalized to the maximum slope value of actin alone for 3 separate experiments in Microsoft Excel and plotted in Prism 10.3.1 (GraphPad Software, Inc).

### TIRF imaging of actin filaments

To make NEM-myosin for immobilizing actin filaments, myosin (Cytoskeleton Inc., MY02) was resuspended in 10 mM imidazole, pH 7 and 0.5 M KCl. NEM (Sigma, E3876) was added at >10-fold molar excess to myosin and allowed to react at room temperature for 1 h before dialyzing overnight at 4°C into 2x HS-TBS (100 mM Tris-HCl, pH 7.5, 1.2 M KCl). Glycerol was added to 50%, sodium azide to 0.02%, and aliquots were stored at −20°C. Actin for TIRF studies was prepared as described above.

Flow cells were prepared using 22×22 mm n=1.5 coverslips (VWR, 48366-227), which were washed for 15 min 0.02% Hellmanex III (Hellma Analytics), 3x with MilliQ water, 15 min in 0.5 M NaOH, 3x with MilliQ water, then dried with a light stream of air. Double-sided Scotch tape was placed on ethanol cleaned microscope slides to create 2 chambers. Cleaned coverslips were placed on top of the tape and pressed with a pipette tip to seal. A solution of 10 nM NEM-myosin in HS-TBS (50 mM Tris-HCl, pH 7.5, 600 mM NaCl) was flowed into chambers for 1 m, then chambers were washed twice with HS-BSA (50 mM Tris-HCl, pH 7.5, 600 mM NaCl, 1% BSA), twice with 2x LS-BSA (50 mM Tris-HCl, pH 7.5, 150 mM NaCl, 1% BSA), and once with 1x TIRF buffer (10 mM imidazole, pH 7, 50 mM KCl, 1 mM MgCl_2_, 1 mM EGTA, 50 mM DTT, 0.2 mM ATP, 50 mM CaCl_2_, 15 mM glucose, 20 µg/mL catalase, 100 µg/mL glucose oxidase, 0.4% methylcellulose).

For TIRF reactions, a solution of 2 µM actin monomers (33% rhodamine-labeled) was supplemented with MgCl_2_ to 50 µM and EDTA to 0.2 mM for 5 min to exchange Ca^2+^ for Mg^2+^. When relevant, 0.5 or 5 nM Sca2/6-AF647 and 3 µM profilin (purified as previously described; (Fedorov et al., 1994)), or an equivalent volume of Buffer A was added. 5 min before imaging, 2x TIRF buffer was added at a 1:1 ratio to the actin and proteins of interest and the mix was flowed into the flow cell.

Imaging was carried out on a Nikon Ti Eclipse Ti2 inverted microscope equipped with a Nikon 60x/1.49 NA CFI Apo TIRF objective and temperature chamber set to 25°C (Oko labs). Single-plane TIRF images were illuminated with a LUNF 4-line laser launch and iLas2 TIRF/FRAP module (Gataca Systems) and captured with an Orca Fusion Gen III sCMOS camera (Hamamatsu) every 5 s for 15-20 m with NIS Elements software (Nikon). For each reaction, the growth of 5 actin filaments in 3 separate movies were measured. Elongation rate was determined over the course of 300 s by manually tracing the barbed ends (fast growing end with visually stable attachment point) of each filament 60 frames apart in ImageJ2/Fiji (v2.9.0/1.53t) and graphed in Prism 10.3.1 (GraphPad Software, Inc).

### Protein alignments, motif identification, and structure prediction

Protein alignments and trees were generated using Geneious Prime 2025.0.3 (https://www.geneious.com). *Rickettsia* RickA sequences were aligned using the Geneious global alignment MUSCLE alignment algorithms. Post alignment, isolated amino acids in RickA were shifted to group with the most relevant neighboring cluster of amino acids. *R. parkeri* strain Portsmouth and *R. bellii* strain RML 369-C RickA and Sca2/6 sequences were submitted to InterPro for sequence motif analysis (Blum et al., 2024). Protein sequences were submitted to the AlphaFold3 server for structure prediction (Abramson et al., 2024).

### SEC-SAXS measurements and analysis

SEC-SAXS studies were performed at the SIBYLS beamline 12.3.1 at the Advanced Light Source (Rosenberg et al., 2022; Classen et al., 2013). 60 µL of 3-8 mg/mL MBP-Sca2/6 or MBP-Sca2/6-L101A/K102A was suspended in SEC running buffer (5 mM Tris, pH 7.5, 1 mM EGTA, 1 mM EDTA, 0.2 mM ATP, 0.5 mM DTT, 150 mM KCl, 1% glycerol). An equimolar amount of actin and 10x molar latrunculin A (Sigma, 428021) were included when indicated. The X-ray wavelength was set to 1.24 Å and the sample-to-detector distance to 2080 mm, generating scattering vectors (q = 4πsinθ/λ, where 2θ is the scattering angle) ranging from 0.01 Å^-1^ to 0.45 Å^-1^. Samples were run over a 1260 Infinity HPLC system (Agilent) connected to a Shodex 803 column equilibrated with SEC running buffer. The eluent was subject to multi-angle light scattering (MALS) and the SAXS.

The radius of gyration (Rg) was calculated for each of the subtracted frames using the Guinier approximation: I(q) = I(0) exp(−q^2^ Rg^2/3^) with the limits qRg < 1.3. The elution peak was compared to the integral of ratios to background and Rg relative to the recorded frame using the program RAW (Hopkins et al., 2017). Final merged SAXS profiles, derived by integrating multiple frames at the elution peak, were used for further analyses. The Guinier plot and pair distribution function (P(r)) were respectively calculated to determine the volume of correlation (Vc) (Rambo and Tainer, 2013) and the maximal inter-particle dimension (Svergun, 1992).

Protein models were generated using Alphafold 3 (Abramson et al., 2024). The conformational variability of proteins was performed by BilboMD restrained by PAE values from Alphafold 3 with low values PAE (< 0.2) grouped as rigid bodies (Pelikan et al., 2009). The generated models were fitted to the SAXS curves using FoXS (Schneidman-Duhovny et al., 2013). The 2-state ensemble model was selected by MultiFoXS (Schneidman-Duhovny et al., 2016). Regions with low PAE (< 0.2) were grouped together as rigid bodies in the const.inp files. Residues 70-180 of Sca2 were bound to actin as a rigid domain based on the PAE matrices.

### Fixed cell imaging

For fixed-cell imaging of cells infected with *R. parkeri* or *R. bellii*, bacteria (WT, pRAM18dRGA, or pRAM18dR-2xCoTagBFP) from 30% prep frozen stocks were added to tissue culture media at an MOI of 0.6-10 (based on PFU measurements described above), aliquoted onto coverslips pre-seeded with 2 x 10^5^ A549 or HMEC-1 cells, and centrifuged at 300 x g for 5 min at room temperature to facilitate infection. Infected cells were incubated at 33°C for 1 h, washed 2x with PBS (Gibco, 10010072) to remove excess bacteria, and tissue culture media was replaced. Infections were allowed to proceed for 15 m - 24 h at 33°C before cells were fixed with 4% paraformaldehyde (Ted Pella Inc, 18505) in PBS for 10 min at room temperature. For fixed-cell imaging of cells infected with *L. monocytogenes,* the cells were diluted into the relevant tissue culture media at an MOI of 5 (based on OD_600_ of 1.0 = 1×10^9^ CFU/ml *L. monocytogenes*; (Ghosh and Higgins, 2018)), and incubated at 37°C for 15 min.

A549 cells were infected as described above, and infections were allowed to proceed at for a total of 8 h before they were fixed as described above. For Arp2/3 inhibitor studies, coverslips were infected with *R. bellii* for 8 h and 24 h or *L. monocytogenes* for 8 h, as described above. At 1 h prior to fixation, coverslips were treated with 200 µM CK689 (Sigma, 182517), CK666 (Sigma, 182515), or CK869 (Sigma, C9124) dissolved in DMSO.

After fixation, coverslips were permeabilized with 0.5% Triton-X 100 (Sigma, T9284) in PBS (Gibco, 10010-023) for 5 min. For localization of surface RickA or Sca2 orthologs, coverslips were blocked for 1 h with 2% BSA (Sigma, A4503) in PBS and then incubated with 1:1000 affinity purified anti-RbSca2/6, anti-RpSca2, or anti-RbRickA in PBS with 2% BSA for 1 h at room temperature. For localization of internal (total) RickA, after blocking, bacteria were permeabilized with lysozyme buffer (PBS, 50 mM glucose, 5 mM EDTA, 5 mg/ml lysozyme (Sigma), and 0.1% Triton X-100) in a humidified chamber for 20 min at 37°C. For quantifying bacteria in Figure 4, coverslips were incubated with 1:500 rabbit anti-I7 (obtained from Ted Hackstadt at the NIH/NIAID Rocky Mountain Laboratories) in PBS with 2% BSA for 30 min at room temperature. All coverslips were incubated with 1:500 568-phalloidin (Molecular Probes, A12380) and 1:500 anti-rabbit 488 (Invitrogen, A-11008) in PBS with 2% BSA for 30 min at room temperature. Coverslips were mounted in ProLong Gold Antifade (Invitrogen, P36930), allowed to cure overnight, and sealed with clear nail polish the next day.

Standard confocal imaging of bacteria, RickA, Sca2/6, and actin was carried out using a Nikon Ti Eclipse microscope equipped with a Yokogawa CSU-XI spinning disc confocal using the 100x VC (1.4 NA) Plan Apo objective and a Clara Interline CCD Camera (Andor Technology). Bacteria were used to localize regions of interest for imaging and 3.5-4.5 µm image stacks were taken with a step size of 0.15 µm to capture all actin tails present in the field of view. Images were imported into ImageJ2/Fiji (v2.9.0/1.53t) (Schindelin et al., 2012) and Z Projection with max intensity was used to collapse the stack. Bacteria, actin tails, and RickA or Sca2/6 puncta were manually quantified and plotted in GraphPad Prism 10.3.1 (GraphPad Software, Inc.).

Lattice structured illumination microscopy images of Sca2 ortholog localization on *R. parkeri* and *R. bellii* were collected at the Rausser College of Natural Resources Biological Imaging Facility at the University of California, Berkeley, using a Zeiss Elyra 7 Lattice Structured Illumination Microscope system running Zen Black using the 63x/1.4NA objective and the system SCMOS cameras. Bacteria with actin tails were used to localize regions of interest for imaging 3-5.6 µm image stacks with a step size of 0.091 µm. Raw 3D images stacks were processed using the SIM function of Zen Black. SIM files were imported into Huygens Professional (Scientific Volume Imaging, v24.10) and image restoration (deconvolution) was performed using CMLE to 0.01% confidence.

To quantify and compare Sca2 ortholog localization and coverage of the *R. parkeri* versus *R. bellii* surface, deconvolved images stacks were analyzed using Imaris 10.2 (Oxford Instruments). Separate surfaces were created for bacteria and Sca2 ortholog to generate area (sum of triangle surfaces for Sca2 ortholog), sphericity (surface area of a sphere with the same volume as the Sca2 ortholog surface/surface area of the Sca2 ortholog surface) and overlapped volume (volume of the overlap between the bacteria and Sca2 ortholog defined surfaces) values used for quantifying Sca2 ortholog coverage of *Rickettsia*. Results were plotted in GraphPad Prism 10.3.1 (Graphpad Software, Inc.). For actin tail intensity line scans, deconvolved images were imported into ImageJ2/Fiji (v2.9.0/1.53t) and a max intensity projection was made for the actin channel of each image. A 1.25 µm line segment was drawn perpendicular to the actin tail 0.5 µm from the base of the bacterium and the intensity values across the perpendicular line segment were saved for one actin tail per image. The intensity values were normalized to the smallest and largest value across individual actin tails and plotted in GraphPad Prism 10.3.1. The *R. parkeri* actin tail width was calculated from the distance between the two peak fluorescence values.

### Live cell imaging

For live cell imaging, 1 x 10^6^ A549 or HMEC-1 cells stably expressing RFP-F-tractin (generated as described above) were plated in a MatTek dish (MatTek, P35G-1.5-20-C) and incubated at 37°C overnight. Mammalian cells were infected with *R. parkeri* pRAM18dRGA or *R. bellii* pRAM18dRGA at an MOI of 1, as described above. Immediately before imaging, media was changed to live cell imaging media (Live Cell Imaging Solution (Molecular Probes, A59688DJ), 10% FBS (ATLAS Biologics, FP-0500-A), 10 mM glucose (Fisher, D16), and 0.01% oxyrase (v/v) (Oxyrase Inc, OF-0005)). Dishes were imaged in an environmental chamber set to 33°C on the Nikon Ti Eclipse microscope described above. Images were taken at 5 s intervals for 5 min in cells where actin tails were clearly observed.

Images were imported into ImageJ2/Fiji (v2.9.0/1.53t) (Schindelin et al., 2012). To calculate motility speed, the Manual Tracking Plug-In was used to track individual bacteria with actin tails over the course of 1 m in 5 s intervals, with the total distance traveled in µm representing the speed µm/min. Motility efficiency was calculated by recording the X and Y pixel coordinates from t = 0 m and t = 1 m timepoints and then using the distance formula in Microsoft Excel to determine the straight-line distance in µm between the 0 m and 1 m coordinates. This value was divided by the total distance traveled over 1 m to obtain a motility efficiency value. Results were plotted in GraphPad Prism 10.3.1 (Graphpad Software, Inc.).

## Acknowledgements

We thank Erin Goley for providing the pRAM18dR-2xCoTagBFP plasmid for using in making fluorescent *R. bellii* and *R. parkeri*. Thank you to Chris Jeans at the UC Berkeley QB3 MacroLab for helping with protein expression and purification of MBP-Sca2 proteins. We thank Denise Schichnes at the UC Berkeley RCNR Biological Imaging Facility and Samantha Smith for training and assistance with LSIM and TIRF microscopy, and Lucy Brennan for advice about protein purification. SAXS data was collected at the Advanced Light Source (ALS) at the SIBYLS beamline, a national user facility operated by Lawrence Berkeley National Laboratory on behalf of the Department of Energy, Office of Basic Energy Sciences, through the Integrated Diffraction Analysis Technologies (IDAT) program, supported by DOE Office of Biological and Environmental Research. Additional support comes from the NIH/NIGMS project ALS-ENABLE, P30 GM124169. This work was supported by NIH/NIAID grant R01 AI109044 to M.D.W. M.C.B. was funded by a Berkeley Fellowship for Graduate Study, NIH/NIGMS training grant T32 GM007232, and a Kathleen L. Miller Fellowship from the Center for Emerging and Neglected Diseases.

## References

Abramson, J., J. Adler, J. Dunger, R. Evans, T. Green, A. Pritzel, O. Ronneberger, L. Willmore, A.J. Ballard, J. Bambrick, S.W. Bodenstein, D.A. Evans, C.-C. Hung, M. O’Neill, D. Reiman, K. Tunyasuvunakool, Z. Wu, A. Žemgulytė, E. Arvaniti, C. Beattie, O. Bertolli, A. Bridgland, A. Cherepanov, M. Congreve, A.I. Cowen-Rivers, A. Cowie, M. Figurnov, F.B. Fuchs, H. Gladman, R. Jain, Y.A. Khan, C.M.R. Low, K. Perlin, A. Potapenko, P. Savy, S. Singh, A. Stecula, A. Thillaisundaram, C. Tong, S. Yakneen, E.D. Zhong, M. Zielinski, A. Žídek, V. Bapst, P. Kohli, M. Jaderberg, D. Hassabis, and J.M. Jumper. 2024. Accurate structure prediction of biomolecular interactions with AlphaFold 3. Nature. 630:493–500. doi:10.1038/s41586-024-07487-w.

Alqassim, S.S. 2022. Functional Mimicry of Eukaryotic Actin Assembly by Pathogen Effector Proteins. Int. J. Mol. Sci. 23:11606. doi:10.3390/ijms231911606.

Alqassim, S.S., I.-G. Lee, and R. Dominguez. 2019. *Rickettsia* Sca2 Recruits Two Actin Subunits for Nucleation but Lacks WH2 Domains. Biophys. J. 116:540–550. doi:10.1016/j.bpj.2018.12.009.

Avvaru, B.S., J. Pernier, and M.-F. Carlier. 2015. Dimeric WH2 repeats of VopF sequester actin monomers into non-nucleating linear string conformations: An X-ray scattering study. J. Struct. Biol. 190:192–199. doi:10.1016/j.jsb.2015.03.008.

Benanti, E.L., C.M. Nguyen, and M.D. Welch. 2015. Virulent *Burkholderia* Species Mimic Host Actin Polymerases to Drive Actin-Based Motility. Cell. 161:348–360. doi:10.1016/j.cell.2015.02.044.

Blanc, G., M. Ngwamidiba, H. Ogata, P.-E. Fournier, J.-M. Claverie, and D. Raoult. 2005. Molecular Evolution of *Rickettsia* Surface Antigens: Evidence of Positive Selection. Mol. Biol. Evol. 22:2073–2083. doi:10.1093/molbev/msi199.

Blanc, G., H. Ogata, C. Robert, S. Audic, K. Suhre, G. Vestris, J.-M. Claverie, and D. Raoult. 2007. Reductive Genome Evolution from the Mother of *Rickettsia*. PLoS Genet. 3:e14. doi:10.1371/journal.pgen.0030014.

Blum, M., A. Andreeva, L.C. Florentino, S.R. Chuguransky, T. Grego, E. Hobbs, B.L. Pinto, A. Orr, T. Paysan-Lafosse, I. Ponamareva, G.A. Salazar, N. Bordin, P. Bork, A. Bridge, L. Colwell, J. Gough, D.H. Haft, I. Letunic, F. Llinares-López, A. Marchler-Bauer, L. Meng-Papaxanthos, H. Mi, D.A. Natale, C.A. Orengo, A.P. Pandurangan, D. Piovesan, C. Rivoire, C.J.A. Sigrist, N. Thanki, F. Thibaud-Nissen, P.D. Thomas, S.C.E. Tosatto, C.H. Wu, and A. Bateman. 2024. InterPro: the protein sequence classification resource in 2025. Nucleic Acids Res. 53:D444–D456. doi:10.1093/nar/gkae1082.

Breitsprecher, D., and B.L. Goode. 2013. Formins at a glance. J. Cell Sci. 126:1–7. doi:10.1242/jcs.107250.

Burke, T.A., A.J. Harker, R. Dominguez, and D.R. Kovar. 2017. The bacterial virulence factors VopL and VopF nucleate actin from the pointed end. J. Cell Biol. 216:1267–1276. doi:10.1083/jcb.201608104.

Burke, T.P., P. Engström, C.J. Tran, I.M. Langohr, D.R. Glasner, D.A. Espinosa, E. Harris, and M.D. Welch. 2021. Interferon receptor-deficient mice are susceptible to eschar-associated rickettsiosis. eLife. 10:e67029. doi:10.7554/elife.67029.

Burkhardt, N.Y., G.D. Baldridge, P.C. Williamson, P.M. Billingsley, C.C. Heu, R.F. Felsheim, T.J. Kurtti, and U.G. Munderloh. 2011. Development of Shuttle Vectors for Transformation of Diverse *Rickettsia* Species. PLoS ONE. 6:e29511. doi:10.1371/journal.pone.0029511.

Cao, L., S. Huang, A. Basant, M. Mladenov, and M. Way. 2024. CK-666 and CK-869 differentially inhibit Arp2/3 iso-complexes. EMBO Rep. 25:3221–3239. doi:10.1038/s44319-024-00201-x.

Carman, P.J., G. Rebowski, R. Dominguez, and S.S. Alqassim. 2023. Single particle cryo-EM analysis of *Rickettsia conorii* Sca2 reveals a formin-like core. J. Struct. Biol. 215:107960. doi:10.1016/j.jsb.2023.107960.

Classen, S., G.L. Hura, J.M. Holton, R.P. Rambo, I. Rodic, P.J. McGuire, K. Dyer, M. Hammel, G. Meigs, K.A. Frankel, and J.A. Tainer. 2013. Implementation and performance of SIBYLS: a dual endstation small-angle X-ray scattering and macromolecular crystallography beamline at the Advanced Light Source. J. Appl. Crystallogr. 46:1–13. doi:10.1107/s0021889812048698.

Colonne, P.M., C.G. Winchell, and D.E. Voth. 2016. Hijacking Host Cell Highways: Manipulation of the Host Actin Cytoskeleton by Obligate Intracellular Bacterial Pathogens. Front. Cell. Infect. Microbiol. 6:107. doi:10.3389/fcimb.2016.00107.

Dominguez, R. 2016. The WH2 Domain and Actin Nucleation: Necessary but Insufficient. Trends Biochem. Sci. 41:478–490. doi:10.1016/j.tibs.2016.03.004.

Dowd, G.C., R. Mortuza, and K. Ireton. 2021. Molecular Mechanisms of Intercellular Dissemination of Bacterial Pathogens. Trends Microbiol. 29:127–141. doi:10.1016/j.tim.2020.06.008.

Ducka, A.M., P. Joel, G.M. Popowicz, K.M. Trybus, M. Schleicher, A.A. Noegel, R. Huber, T.A. Holak, and T. Sitar. 2010. Structures of actin-bound Wiskott-Aldrich syndrome protein homology 2 (WH2) domains of Spire and the implication for filament nucleation. Proc. Natl. Acad. Sci. 107:11757–11762. doi:10.1073/pnas.1005347107.

Fan, E., N. Chauhan, D.B.R.K.G. Udatha, J.C. Leo, and D. Linke. 2016. Type V Secretion Systems in Bacteria. Microbiol. Spectr. 4:10.1128/microbiolspec.vmbf-0009–2015. doi:10.1128/microbiolspec.vmbf-0009-2015.

Fedorov, A.A., T.D. Pollard, and S.C. Almo. 1994. Purification, Characterization and Crystallization of Human Platelet Profilin Expressed in *Escherichia coli*. J. Mol. Biol. 241:480–482. doi:10.1006/jmbi.1994.1522.

Figueroa-Cuilan, W.M., O. Irazoki, M. Feeley, E. Smith, T. Nguyen, F. Cava, and E.D. Goley. 2023. Quantitative analysis of morphogenesis and growth dynamics in an obligate intracellular bacterium. Mol. Biol. Cell. 34:ar69. doi:10.1091/mbc.e23-01-0023.

Gautreau, A.M., F.E. Fregoso, G. Simanov, and R. Dominguez. 2022. Nucleation, stabilization, and disassembly of branched actin networks. Trends Cell Biol. 32:421–432. doi:10.1016/j.tcb.2021.10.006.

Ghosh, P., and D.E. Higgins. 2018. *Listeria monocytogenes* Infection of the Brain. J. Vis. Exp. doi:10.3791/58723.

Gouin, E., C. Egile, P. Dehoux, V. Villiers, J. Adams, F. Gertler, R. Li, and P. Cossart. 2004. The RickA protein of *Rickettsia conorii* activates the Arp2/3 complex. Nature. 427:457–461. doi:10.1038/nature02318.

Gouin, E., H. Gantelet, C. Egile, I. Lasa, H. Ohayon, V. Villiers, P. Gounon, P.J. Sansonetti, and P. Cossart. 1999. A comparative study of the actin-based motilities of the pathogenic bacteria *Listeria monocytogenes, Shigella flexneri* and *Rickettsia conorii*. J. Cell Sci. 112:1697–1708. doi:10.1242/jcs.112.11.1697.

Haglund, C.M., J.E. Choe, C.T. Skau, D.R. Kovar, and M.D. Welch. 2010. *Rickettsia* Sca2 is a bacterial formin-like mediator of actin-based motility. Nat. Cell Biol. 12:1057–1063. doi:10.1038/ncb2109.

Harris, E.K., K. Jirakanwisal, V.I. Verhoeve, C. Fongsaran, C. Suwanbongkot, M.D. Welch, and K.R. Macaluso. 2018. Role of Sca2 and RickA in the Dissemination of *Rickettsia* parkeri in Amblyomma maculatum. Infect. Immun. 86:10.1128/iai.00123-18. doi:10.1128/iai.00123-18.

Heinzen, R.A., S.F. Hayes, M.G. Peacock, and T. Hackstadt. 1993. Directional actin polymerization associated with spotted fever group *Rickettsia* infection of Vero cells. Infect. Immun. 61:1926–1935. doi:10.1128/iai.61.5.1926-1935.1993.

Hopkins, J.B., R.E. Gillilan, and S. Skou. 2017. BioXTAS RAW: improvements to a free open-source program for small-angle X-ray scattering data reduction and analysis. J. Appl. Crystallogr. 50:1545–1553. doi:10.1107/s1600576717011438.

Jeng, R.L., E.D. Goley, J.A. D’Alessio, O.Y. Chaga, T.M. Svitkina, G.G. Borisy, R.A. Heinzen, and M.D. Welch. 2004. A *Rickettsia* WASP-like protein activates the Arp2/3 complex and mediates actin-based motility. Cell. Microbiol. 6:761–769. doi:10.1111/j.1462-5822.2004.00402.x.

Johnson, H.W., and M.J. Schell. 2009. Neuronal IP3 3-Kinase is an F-actin–bundling Protein: Role in Dendritic Targeting and Regulation of Spine Morphology. Mol. Biol. Cell. 20:5166– 5180. doi:10.1091/mbc.e09-01-0083.

Karkouri, K.E., E. Ghigo, D. Raoult, and P.-E. Fournier. 2022. Genomic evolution and adaptation of arthropod-associated *Rickettsia*. Sci. Rep. 12:3807. doi:10.1038/s41598-022-07725-z.

Kleba, B., T.R. Clark, E.I. Lutter, D.W. Ellison, and T. Hackstadt. 2010. Disruption of the *Rickettsia rickettsii* Sca2 Autotransporter Inhibits Actin-Based Motility. Infect. Immun. 78:2240–2247. doi:10.1128/iai.00100-10.

Kudryashova, E., Ankita, H. Ulrichs, S. Shekhar, and D.S. Kudryashov. 2022. Pointed-end processive elongation of actin filaments by *Vibrio* effectors VopF and VopL. Sci. Adv. 8:eadc9239. doi:10.1126/sciadv.adc9239.

Lamason, R.L., E. Bastounis, N.M. Kafai, R. Serrano, J.C. del Álamo, J.A. Theriot, and M.D. Welch. 2016. *Rickettsia* Sca4 Reduces Vinculin-Mediated Intercellular Tension to Promote Spread. Cell. 167:670–683.e10. doi:10.1016/j.cell.2016.09.023.

Lamason, R.L., and M.D. Welch. 2017. Actin-based motility and cell-to-cell spread of bacterial pathogens. Curr. Opin. Microbiol. 35:48–57. doi:10.1016/j.mib.2016.11.007.

Lehman, S.S., V.I. Verhoeve, T.P. Driscoll, J.F. Beckmann, and J.J. Gillespie. 2024. Metagenome diversity illuminates the origins of pathogen effectors. mBio. 15:e00759–23. doi:10.1128/mbio.00759-23.

Liverman, A.D.B., H.-C. Cheng, J.E. Trosky, D.W. Leung, M.L. Yarbrough, D.L. Burdette, M.K. Rosen, and K. Orth. 2007. Arp2/3-independent assembly of actin by *Vibrio* type III effector VopL. Proc. Natl. Acad. Sci. 104:17117–17122. doi:10.1073/pnas.0703196104.

Madasu, Y., C. Suarez, D.J. Kast, D.R. Kovar, and R. Dominguez. 2013. *Rickettsia* Sca2 has evolved formin-like activity through a different molecular mechanism. Proc. Natl. Acad. Sci. 110:E2677–E2686. doi:10.1073/pnas.1307235110.

Namgoong, S., M. Boczkowska, M.J. Glista, J.D. Winkelman, G. Rebowski, D.R. Kovar, and R. Dominguez. 2011. Mechanism of actin filament nucleation by *Vibrio* VopL and implications for tandem W domain nucleation. Nat. Struct. Mol. Biol. 18:1060–1067. doi:10.1038/nsmb.2109.

Ngwamidiba, M., G. Blanc, H. Ogata, D. Raoult, and P.-E. Fournier. 2005. Phylogenetic Study of *Rickettsia* Species Using Sequences of the Autotransporter Protein-Encoding Gene sca2. Ann. N. York Acad. Sci. 1063:94–99. doi:10.1196/annals.1355.015.

Nolen, B.J., N. Tomasevic, A. Russell, D.W. Pierce, Z. Jia, C.D. McCormick, J. Hartman, R. Sakowicz, and T.D. Pollard. 2009. Characterization of two classes of small molecule inhibitors of Arp2/3 complex. Nature. 460:1031–1034. doi:10.1038/nature08231.

Pan, Y.-S., X.-M. Cui, L.-F. Du, L.-Y. Xia, C.-H. Du, L. Bell-Sakyi, M.-Z. Zhang, D.-Y. Zhu, Y. Dong, W. Wei, L. Zhao, Y. Sun, Q.-Y. Lv, R.-Z. Ye, Z.-H. He, Q. Wang, L.-J. Li, M.-G. Yao, T. Xiong, J.-F. Jiang, W.-C. Cao, and N. Jia. 2022. Coinfection of Two *Rickettsia* Species in a Single Tick Species Provides New Insight into *Rickettsia-Rickettsia* and *Rickettsia*-Vector Interactions. Microbiol. Spectr. 10:e02323–22. doi:10.1128/spectrum.02323-22.

Pelikan, M., G. Hura, and M. Hammel. 2009. Structure and flexibility within proteins as identified through small angle X-ray scattering. Gen. Physiol. Biophys. 28:174–189. doi:10.4149/gpb_2009_02_174.

Rambo, R.P., and J.A. Tainer. 2013. Accurate assessment of mass, models and resolution by small-angle scattering. Nature. 496:477–481. doi:10.1038/nature12070.

Reed, S.C.O., R.L. Lamason, V.I. Risca, E. Abernathy, and M.D. Welch. 2014. *Rickettsia* Actin-Based Motility Occurs in Distinct Phases Mediated by Different Actin Nucleators. Curr. Biol. 24:98–103. doi:10.1016/j.cub.2013.11.025.

Rosenberg, D.J., G.L. Hura, and M. Hammel. 2022. Size exclusion chromatography coupled small angle X-ray scattering with tandem multiangle light scattering at the SIBYLS beamline. Methods Enzym. 677:191–219. doi:10.1016/bs.mie.2022.08.031.

Rymaszewska, A., and M. Piotrowski. 2024. Rickettsia Species: Genetic Variability, Vectors, and Rickettsiosis—A Review. Pathogens. 13:661. doi:10.3390/pathogens13080661.

Schindelin, J., I. Arganda-Carreras, E. Frise, V. Kaynig, M. Longair, T. Pietzsch, S. Preibisch, C. Rueden, S. Saalfeld, B. Schmid, J.-Y. Tinevez, D.J. White, V. Hartenstein, K. Eliceiri, P. Tomancak, and A. Cardona. 2012. Fiji: an open-source platform for biological-image analysis. Nat. Methods. 9:676–682. doi:10.1038/nmeth.2019.

Schneidman-Duhovny, D., M. Hammel, J.A. Tainer, and A. Sali. 2013. Accurate SAXS Profile Computation and its Assessment by Contrast Variation Experiments. Biophys. J. 105:962– 974. doi:10.1016/j.bpj.2013.07.020.

Schneidman-Duhovny, D., M. Hammel, J.A. Tainer, and A. Sali. 2016. FoXS, FoXSDock and MultiFoXS: Single-state and multi-state structural modeling of proteins and their complexes based on SAXS profiles. Nucleic Acids Res. 44:W424–W429. doi:10.1093/nar/gkw389.

Scott, A.T., C.J. Vondrak, A.G. Sanderlin, and R.L. Lamason. 2022. Rickettsia parkeri. Trends Microbiol. 30:511–512. doi:10.1016/j.tim.2022.01.001.

Sears, K.T., S.M. Ceraul, J.J. Gillespie, E.D. Allen, V.L. Popov, N.C. Ammerman, M.S. Rahman, and A.F. Azad. 2012. Surface Proteome Analysis and Characterization of Surface Cell Antigen (Sca) or Autotransporter Family of *Rickettsia typhi*. PLoS Pathog. 8:e1002856. doi:10.1371/journal.ppat.1002856.

Serio, A.W., R.L. Jeng, C.M. Haglund, S.C. Reed, and M.D. Welch. 2010. Defining a Core Set of Actin Cytoskeletal Proteins Critical for Actin-Based Motility of *Rickettsia*. Cell Host Microbe. 7:388–398. doi:10.1016/j.chom.2010.04.008.

Shaner, N.C., M.Z. Lin, M.R. McKeown, P.A. Steinbach, K.L. Hazelwood, M.W. Davidson, and R.Y. Tsien. 2008. Improving the photostability of bright monomeric orange and red fluorescent proteins. Nat. Methods. 5:545–551. doi:10.1038/nmeth.1209.

Shere, K.D., S. Sallustio, A. Manessis, T.G. D’Aversa, and M.B. Goldberg. 1997. Disruption of IcsP, the major *Shigella* protease that cleaves IcsA, accelerates actin-based motility. Mol. Microbiol. 25:451–462. doi:10.1046/j.1365-2958.1997.4681827.x.

Steinhauer, J., R. Agha, T. Pham, A.W. Varga, and M.B. Goldberg. 1999. The unipolar *Shigella* surface protein IcsA is targeted directly to the bacterial old pole: IcsP cleavage of IcsA occurs over the entire bacterial surface. Mol. Microbiol. 32:367–377. doi:10.1046/j.1365-2958.1999.01356.x.

Stradal, T.E.B., and M. Schelhaas. 2018. Actin dynamics in host–pathogen interaction. FEBS Lett. 592:3658–3669. doi:10.1002/1873-3468.13173.

Svergun, D.I. 1992. Determination of the regularization parameter in indirect-transform methods using perceptual criteria. J. Appl. Crystallogr. 25:495–503. doi:10.1107/s0021889892001663.

Tam, V.C., D. Serruto, M. Dziejman, W. Brieher, and J.J. Mekalanos. 2007. A Type III Secretion System in *Vibrio cholerae* Translocates a Formin/Spire Hybrid-like Actin Nucleator to Promote Intestinal Colonization. Cell Host Microbe. 1:95–107. doi:10.1016/j.chom.2007.03.005.

Tanaka, M., and H. Shibata. 1985. Poly(l-proline)-binding proteins from chick embryos are a profilin and a profilactin. Eur. J. Biochem. 151:291–297. doi:10.1111/j.1432-1033.1985.tb09099.x.

Teysseire, N., C. Chiche-Portiche, and D. Raoult. 1992. Intracellular movements of *Rickettsia conorii* and *R. typhi* based on actin polymerization. Res. Microbiol. 143:821–829. doi:10.1016/0923-2508(92)90069-z.

Tran, C.J., Z. Zubair-Nizami, I.M. Langohr, and M.D. Welch. 2025. The *Rickettsia* actin-based motility effectors RickA and Sca2 contribute differently to cell-to-cell spread and pathogenicity. mBio. 16:e02563–24. doi:10.1128/mbio.02563-24.

Valencia, D.A., and M.E. Quinlan. 2021. Formins. Curr. Biol. 31:R517–R522. doi:10.1016/j.cub.2021.02.047.

Van Kirk, L.S., S.F. Hayes, and R.A. Heinzen. 2000. Ultrastructure of *Rickettsia rickettsii* Actin Tails and Localization of Cytoskeletal Proteins. Infect. Immun. 68:4706–4713. doi:10.1128/iai.68.8.4706-4713.2000.

Voss, O.H., and M.S. Rahman. 2021. *Rickettsia*-host interaction: strategies of intracytosolic host colonization. Pathog. Dis. 79:ftab015. doi:10.1093/femspd/ftab015.

Welch, M.D., A. Iwamatsu, and T.J. Mitchison. 1997. Actin polymerization is induced by Arp 2/3 protein complex at the surface of *Listeria monocytogenes*. Nature. 385:265–269. doi:10.1038/385265a0.

Yu, B., H.-C. Cheng, C.A. Brautigam, D.R. Tomchick, and M.K. Rosen. 2011. Mechanism of actin filament nucleation by the bacterial effector VopL. Nat. Struct. Mol. Biol. 18:1068–1074. doi:10.1038/nsmb.2110.

Zahm, J.A., S.B. Padrick, Z. Chen, C.W. Pak, A.A. Yunus, L. Henry, D.R. Tomchick, Z. Chen, and M.K. Rosen. 2013. The Bacterial Effector VopL Organizes Actin into Filament-like Structures. Cell. 155:423–434. doi:10.1016/j.cell.2013.09.019.

Zhang, J. 2003. Evolution by gene duplication: an update. Trends Ecol. Evol. 18:292–298. doi:10.1016/s0169-5347(03)00033-8.

