## Supplemental Figures and Movie Legends for "Divergent *Rickettsia* species exhibit distinct mechanisms of actin-based motility"

### A RickA

```
R. rickettsii M---VKEIDINKLLAQENNA LNAILSHVNELCKQNKQLQG LIEIQNETKELEKEHNRSLP WFKRFVKTVSNVKYILIKSE  77
R. parkeri    M---VKEIDINKLLAQENNA LNTILSQVNELCKQNKQLQG LIEIQNETKQLEKEHNRSLP WFKRFVKTVSNVKYILIKSE  77
R. akari      M---AQEIDINKLLAQENNA LNAILSQVNELCTQNKKLQG LIEIQNKTKELAKEHNRSLP WFKRLVKTVSNVKYIFVKSE  77
R. felis      MYGVAKEIDINKLLAQENNA LNTILSQVNELCEQNKKLQG LIEIQNETKELEKEHNRSLP WFKRLVKTVSNVKYIFVKSE  80
R. bellii     M---AKITELDHHLNQEKEA LDKVVSNLNELCEHNQKLQG FIEIQKEVKELKKEHIKSLS WFKKLINTVSNIKYVFVKSE  77

R. rickettsii EQLTNEAIKYNNKILKDIDN KIYNIAEKSAPLKQALQEEI EKSFKDLTKKDLSKDQRARL SEVFFS-YKSKPERFSALHM  156
R. parkeri    EQLTNEAIKYNNKILKDIDN KIYNIAEKSAPLKQALQEEI EKNFKDLTKKDLSKDQRARL SEVFFS-YKSKPERFSSLHM  156
R. akari      EQLTNEAIKYNNKVLKSIDN KIYNIAAKSAPLKQELQEEI AKNFKDLTKKDLSKEQRERL SEVYFS-YKSKPERFSALHI  156
R. felis      EQLTNEAIKYNNKILKDIDN KIYNIAEKSAPLKQELQEEI EKNFKDLTKKDLSKEQRERL SEVYFS-YKSKPERFSALNM  159
R. bellii     EQLAKDAIEQNNKLLKRIDN TILSVADKSGPLKQELQKEL RKNFENLAKKDLSKDQRERL SNLLNNEYAANPQKFAQLPM  157

R. rickettsii TNPLQFINAEALEKQFNSLN ATKQNIQNLISENSNIKELK EIQKQVAEIRAEVPHTFFEK LNNIWQNVKNVFVNNSEQVL  236
R. parkeri    TNPLQFINAEALEKQYNSLN ATKQNIQNLISENSNIKELK EIQKQVAEIRAEVPHTFFEK LNNIWQNVKNVFVNNSEQVL  236
R. akari      TNPLQFINAEELEKQYNSLN ATRQNIQNLISENSNIKELK EIQKQVAEIREEIPYTFFEK LNNIWQNVKNVFVNNKEKIL  236
R. felis      TNPLQFIKAEELEKQYNSLN ATKQNIQNLISENSNIKELK EIQKQVAEIREEIPYTFFEK LNNIWQNVKNVFVNNSEQVL  239
R. bellii     SKPLHFPNAEELENQHNDLK VIQQNVLNLLTENSNIEELK KIQKQVAEIREEVPFTKLEK LNNFWQKIKNIFVNNSEQVL  237

R. rickettsii AKNKESNTRTIRKIDEQLYK TQHKFEELIENKERNIKDII AKLPDNEELQKIVSNLTNHM ASKKEPILANVSLAKPLENN  316
R. parkeri    AKNKESNTRTIRKIDEQLYK TKHKFEELIENKERNIKDII AKLPDNEKLQKIVSNLTNHM ASQKEPILANASLAKPLENN  316
R. akari      AKNKESNTRTIRKIDEQLYK TKHKFEELIENKERNINDII SKLPDNEELKNIVSNFANHM TSKKEPILTTSSIAKPLENN  316
R. felis      AKNKESNTRAIRKIDEQLYK TKHKFEELIENKERNINDII AKLPDNEELQKIVSNLANHM TSKKEPILTTSSIAKPLENN  319
R. bellii     AKNKENNTKTIINIEEKLHK ANNKFFELVSNKKQDIENII SNLPDSKRLEAIKEKLQKHI NVKDTNNIAEQASAAQLQSA  317

R. rickettsii I------------------- ------PPPPPPPPPLPDSN IP----PPPPPPLPGNNI-- ----PPPPPPPPPPM--APV  359
R. parkeri    ITPPSPLPENNTPSPPPPPL PENNT---PSPPPPPLPENN TPSPPPPPPPPPLPENNTPS PPPPPPPPPPPPPPM--APA  391
R. akari      ITR----------------- ------------PPLLSQNN IP------SPPPLSKNNI-- ----PPPPPPPPPPM--PTA  353
R. felis      VT------------------ -----------PPPPLTKNN IP---PPPPPPPLSKNNIL- -----PPPPPPMPTM--APA  359
R. bellii     ETKPTAVVLPNNAIPTTPPV TEEKTFTPPPAPPPPMPTDN IPTPLPVSKAEATEHKNVET AASNVPPPPPPPMPTGNVPP  397

R. rickettsii KT--LSKAVEATT-VKKLEN QPRPSIDTSDLMREIAGPKK LKKVE--------------- ------------FDPNTGKP  409
R. parkeri    KT--LSKPVESTT-VKKLEN QPSPSIDTFDLMRELAGPKK LKKVE--------------- ------------FDPNTGKP  441
R. akari      QTDALSKPVGVTT-VKKLEN QQRPSLDTSDLMREIAGPNN LRKVEKTDVKIQDSRDLLLQ SIRGEHKLRKVAFDPNTGKP  432
R. felis      QTETLSKPVGVTTTVKKLEN QPRPSIDTSDLMREIAGPKN LRKVEKTDVKTQDSRDLLLQ SIRGEHKLRKVEFDPNTGKP  439
R. bellii     PPPVGDNTVTSTPQKAKETN QPRPAVDTTNLMKQIQGGFN LKKIE--------------- ------------YGED-GKP  449

R. rickettsii VAHSHSKPAQNVNKPSGLES IFARRVAIEMSDSSS----S ESDSGNWSDVSVNRNKSKML KTKGERDAKMTTHAQKI-NN  484
R. parkeri    VAHPHSKPAQNVNKPSGLES IFARRVAIEVSDSSS----S ESDSGNWSDVSVNRNKSKIL KTKGERDAKMTTHAQKI-NN  516
R. akari      VAHSHSKPAKNVNQPNGVAS ILARRVAMEMSDSSSSGSES DSDSGNWSDISVNSDKPKAL KNRRERDGKRTTHAQKILSN  512
R. felis      VAHSHSKPAQNVSKPNGVAS ILARRVAMEMSDSSSS-SGS ESDSGNWSDASVNSNKPKAL KTRGERDAKTTTHAQKILSN  518
R. bellii     IP-KNKEDTKETSDPI-IAA LNKIRSAKVSSDSERSNSDS GTDSGWASDVST---RSKKV LTRRERNAK----------- 513

R. rickettsii RNSQKPSFVR-                                                                         494
R. parkeri    RNSQKPSFVR-                                                                         526
R. akari      RSSQKPSFVRS                                                                         523
R. felis      RSSQKPSFVRS                                                                         529
R. bellii     QSQQR------                                                                         518
```

### B Sca2

```
Consensus     MNLQNSHSKKYVLTFFMSTC LLTSSFLSTSARAASFKDLV SKTPXWXKHNSTQQQNIWKD LTPNEKIKKWQEAALVPSFT  80
R. rickettsii MNLQNSHSKKYVLTFFMSTC LLTSSFLSTSARAASFKDLV SKTPAWAKHNSTQQQNIWKD LTPTEKIKKWQEAALVPSFT  80
R. parkeri    MNLQKSHSKKYVLTFFMSTC LLTSSFLSTSARAASFKDLV SKTPAWEKHNSTQQQNIWKD LTPNEKIKKWQEAALVPSFT  80
R. akari      MKLQNSYSKKYVLTFFMSTC LFTSSFLSTSTRAASFKDLV SKTPTWQKHNAKQQQNIWKD FTPNEKIKKWQEANLIPSFT  80
R. felis      MSLQNSHSKKYVLTFFMSTC LLTSSFLSTSARAASFTQLA NQIPTLSGLSEVQRKQKWNS YTLQEKQEAWRRAKLTPDFV  80
R. typhi      -------------------- -------------------- -------------------- --------------------
R. bellii     -------------------- -------------------- -------------------- --------------------

Consensus     QAQNDLGIKYKETDLSSFLD NTRHKARQARAEILLYIERI KQQDFDTKKQEYINQGVVPT DIEAATNLGISYDPSKIDNX 160
R. rickettsii QAQNDLGIKYKETDLSSFLD NTRHKARQARAEILLYIERI KQQDFDTKKQAYINQGVVPT DIEAATNLGISYDPSKIDHN 160
R. parkeri    QAQNDLGIKYKETDLSSFLD NTRHKARQARAEILLYIERV KQQDFDTKKQEYINQGVVPT DIEAATNLGISYDPSKIDNK 160
R. akari      QAQDDLGIQYKETDLSSFLD NTRHKARQARAEILLYIQRI KQQDFDTKKQEYIKQGIVPT YMEAATILGISYDPSKIYDN 160
R. felis      QAMIDMQTGFTESDLSSRNN KTRHKAREKKSELDLYIAGT K-QGFKEKVDGYISQGKIPT PEEAAQNLEIDYDAKKTDNK 159
R. typhi      -------------------- -------------------- -------------------- --------------------
R. bellii     -------------------- -------------------- -------------------- --------------------

Consensus     VEXDQXVRRAEKDKKAXIXL YISSINRDIKYKHYVDNDII PEMQEVRTALNMNKDDAQSF VASIRTEIMENAKGQYIADS 240
R. rickettsii VEHDQKVRRAEKDKKAVIEL YISSINRDIKYKHYVDNDII PEMQEVRTALNMNKDDAQSF VASIRTEIMENAKGQYIADS 240
R. parkeri    VEHDQKVRRAEKDKKAVIEL YISSINRGIKYKHYVDNDII PEMQEVRTALNMNKDDAQSF VASIRTEIMENAKGQYIADS 240
R. akari      VEKDQNVRRAEKDKKALIDL YISSIKRDIKYRHYVNNHII PEIKEVKTALNMDKDDAESF VLSIRTEIMENAKEQYIADR 240
R. felis      LEKNQNVRRVEKDKKALLDL YIHSITTSVKEKEYITTGQV PELGELEKALNISKEEAKHR RTIIRDQVMANERPKLVRSG 239
R. typhi      -------------------- -------------------- -------------------- --------------------
R. bellii     -------------------- -------------------- -------------------- --------------------

Consensus     HIPTEKELKKKFGISRDDNR DGYIKSIRLKVMDKEKPQYI ADSHIPTEKELEQKFGXDK- GEATNYIASIATQMMLGKKS 319
R. rickettsii HIPTEKELKKKFGISRDDNR DGYIKSIRLKVMENAKGQYI ADSHIPTEKELEQKFGADK- GEATNYIASIATQMMLGKKS 319
R. parkeri    HIPTEKELKKKFGISRDDNR DGYIKSIRLKVMDKEKPQYI ADSHIPTEKELEQKFGVDK- GEATNYIASIATQMMLGKKS 319
R. akari      HIPTEKELKNRFGISRDDNR DAYIKSIRLKVMDKEKPQYI AANSIPTEKELEQKFGCDK- GEATNYIASIATQKMLNKKA 319
R. felis      T------------------- -------------------- ----VLTKKELHKRFGKDTT TDDTKYIDDITTEVMYTKKQ 276
R. typhi      -------------------- -------------------- -------------------- --------------------
R. bellii     -------------------- -------------------- -------------------- --------------------

Consensus     YYIDNNIIPNXDELMNEFKI GXVKAXSYINQIKAGIEAXQ XLNNNDTT-KPSTGHSQKKS GXKNDXWYMSNQXINXTETS 398
R. rickettsii YYIDNNIIPNTDELMNEFNI GPVKATSYINQIRAGIEAKQ FLNNNDTT-KPSTGHSQKKS GSKNDHWYMSNQSINDTGTS 398
R. parkeri    YYIDNNIIPNADELMNEFKI GPVKATSYINQIKAGIEANQ FLNNNDTT-KPSTGRSQQKS GSKNNPWYMSNQSIHNTETS 398
R. akari      YYIDNNIIPAVEELKQEFRI GKIKANSYIQQITDGINANQ LLNNNNTT-KPSTVGSTKKI ETKSDNWYMSNQGINTTETS 398
R. felis      GYVNTDFLPKISEIMNEFKV DKGRANLYLNQIKAGIEAKL LADNNQTTTKPFTKHSRTTT NTA---------GI---SSG 344
R. typhi      -------------------- -------------------- -------------------- --------------------
R. bellii     -------------------- -------------------- -------------------- --------------------

Consensus     SXIXTGRXKKQRYFFDPIST FKTHFNXKANKGNLTQSQXN IXRIIQQEENIEEFKNLIKT BPIAALLLQ----VXSSYKQ 474
R. rickettsii SRIFTGREKKQRYFFDPIST FKTHFNTKASKGNLTQSQHT IKRIIQQEENIAEFKNLIKT DPIAALTLQ----VGSSYKQ 474
R. parkeri    SQISTGRDKKQRYFFDSIST FKTQFNAKANKGNLTLSQQN INKLIEQEENIEQFKDLIKT NPIAALLLQ----VDSSYKN 474
R. akari      SVVTTGRKEKQSYYFDPIST FKAHFNNKENNGNLTQPQHN INRIIQQEENIEEFKNLIKT DPIAALNLT----VDSSYKK 474
R. felis      VPFDTGRTKPETKSFDFKRS MYSLLN-RKQEDQLSKTEQH LKQQIKLEENKEEFKEILTK NPIDALLFAEQSNLGNSFKQ 423
R. typhi      -------------------- -------------------- -------------------- --------------------
R. bellii     -------------------- -------------------- -------------------- --------------------

Consensus     EAVTXILSDFNDNTIQRVLF XNDXGQLDFKTNIDVKNRPI LQTLLENSSSEEKTKFAERI QDY----ATRNISNSQFEEK 550
R. rickettsii EAVTTILSDFNDNTIQRVLF SNDKGQLDFKTNIDVKNRPI LQELLENSSSEEKTKFAERI QDY----ATRNISHSQFEEK 550
R. parkeri    QAVKIILKDRNDNTIQRLLF TNDTGQLDFNTNIKVKNRPI LQTFLNNSTSKDKTKFAEII QDY----ATRNISNSQFEEK 550
R. akari      EAVTSILSDFNDDTIQRVLF SDDMGQLDFKTNIDVKNRPI LKALLENSSSEEKTKFAERI QDY----ATRNISNSQFEEK 550
R. felis      EAISNIDL---SKDISRILF TVD----------DKGNRTI LNTILTTTPE-HKDELIKQA QHHAIQTLPTSISDKDVSDK 489
R. typhi      -------------------- -------------------- -------------------- --------------------
R. bellii     -------------------- -------------------- -------------------- --------------------

Consensus     ARLDLIKLAASKDKSSVENF LTLQLELKNRMQPYIVNSGY ILTPEIVXEINIELKNXGLI XDSLTKDYMIKLAKEVNXHT 630
R. rickettsii ARLDLIKLAASKDKRSVENF LTLQLELKNRMQPYIVNSAY ILTPEIVKEINIELKNKGLI RDSLTKDYMIKLAKEVNNHT 630
R. parkeri    ARLDLIKIAASKDKSSVENF LTLQLELKNRMQPYIVNSVY ILTPEIVKEINIELKNKGLI RDSLTKDYMIKLAKEVNNHT 630
R. akari      ARLDLIKLAASKDRSLVEKF LALQLELKNKMQSHIVKSEY ILTPKIVAEINIELKNQGLI IDSLTKDNMIKLAKEVNKQA 630
R. felis      KKLTLATLAATEDKKVLEEA LD----------NWLSTNGY K------------------- ----------------RKPE 524
R. typhi      -------------------- -------------------- -------------------- ------------------MT   2
R. bellii     -------------------- ---------------MSLGY KLKSTL-------------- -----LKYSFVVAISINLLA  26
```

```
Consensus    LNSVI-KVILSDXNXLSNET NKILGLAVGNNANNLX-QTQ SGIPNPPP-LPLN--XPPPP PPPX--X----------XXI  693
R. rickettsii LNSVI-KVILSDSNILSNET NKILGLAVGNNANNLE-QTQ SGIPNPPP-LPLN------- -----------------GDI  683
R. parkeri   LNSVI-KVILSDSNILSNEA NKILGLAVGNNANNLE-QTQ SDIPNPPP-LPLN------- -----------------GGI  683
R. akari     LNSAI-KVILSDNNTLSNET NKILGLAVGNNANNLA-QTQ SGMPNPPP-LPLSGGIPNPP PLPLSGGIPNPPPLPLSGGI  707
R. felis     VESLI-SILLSDETTLKAGI DKIFELPVENNVNNNSNKGQ NGTPILPPTPPLNGSMPPSP PPPLLNGT------------  591
R. typhi     ITGKLKSVLLTY-------- SFIILINL---LTAMS---- -------ESLASSWNPPPPP PPPPIEGLNNFKSL------   54
R. bellii    I------------------N SGILLLTL-------NNKAE AAPPPPPPPPPP---PPPPP PPPPP------P------PT   66

Consensus    PNPPPLPLNGSXPPXPXLHS QGFSSNSXHFDLNXLQAEYP HIHSLYIQFXHNTTVQSKAP XQPTXXX-XXX-RSXPETAY  771
R. rickettsii PNPPPLPLNGSMPPPPPLHS QGFSSNSKHFDLNQLQTEYP HIHSLYIQFTHNTTVQSKAP LQPTASSATSTGRSTPETAY  763
R. parkeri   PNPPPLPLNGSIPP-PPLHS QGFISNSQHFDLNQLQTEYP HIHSLYIQFTHNTTVQSKAP LQPTASSTTSTVRSTPETAY  762
R. akari     PNPPPLPLNGSMPPPPHLNS QGFISNSNNLDLNKLQAEYS HIHSLYTQFILNTTVQPKVL PQPTASSATSTERSEPETAY  787
R. felis     ------------PTSTAF-- -NNSNPNHKFDLKNFEATYP RLYKSYNEFIQNTTSASQSQ ATTT-------SNNIPDTKA  649
R. typhi     -NP------------PSKHS LGNSSKVEQSQHTNKNCTTS NI---QMQFS-----QSLEV PT-------------PTGEF  100
R. bellii    PPPPPLPKTPPVDPKSAA-- ------------KDINAELG PTFNPYDTLKKNRKNI---- ---------------K----  109

Consensus    AKLYXEYRTETGGKKANDLQ DQLIKRQADLTNVIRQILTE SYANXGADEKTLVNLFSIST PEIAEKAKEAFNTLAQDXYI  851
R. rickettsii AKLYAEYRTETGGTKANDLQ DQLIKRQADLTNVIRQILTE SYANQGADEKTLVNLFSIST PEIAEKAKEAFNTLAQDQYI  843
R. parkeri   AKLYAEYRTETGGTKANDLQ DQLIKRQADLTNVIRQILTE SYANQGADEKTLLNLFSIST PEIAEKAKEAFNTLAQDQYI  842
R. akari     AKLYVEYRTETGGKKAYDLQ DQLIKRQADLTNVIRQILTT SYANQGADEKTLVNVFSIST PEIEAKAKDVFNKLVQDPYI  867
R. felis     ---------KMG--ESLELL KQKVAKQNEVIGLIHNEVTK LY---NFSPKTFVNLFNTEN EEIIKKIEQIAK----REDI  711
R. typhi     GKKYSKLINPKTSK-----Q NLKIEFPKKIISLVRDELTL KYFT--ADIERILTLFSKKD DLILSEAKEKYKEVIQDPYI  173
R. bellii    -----------NKKDNSDLE AFL-----NLNGI------- ---------AQMSKLFSIVL TQHT------DNDELRARFF  151

Consensus    KDITVNXKKTITSEEIIK-- -----NLFNEDTDDAVKRIL LSSCKISEELKKPIKLXFN- ---QX---------------  905
R. rickettsii KDITVNGKKTITSEEIIK-- -----NLFNEDTDDAVKRIL LSSCKISEELKRPIKLEFN- ---KS---------------  897
R. parkeri   KDITVNGKKTITSEEIIK-- -----NLFNEDTDDAVKRIL LSSCKISEELKRPIKLKFN- ---QS---------------  896
R. akari     QYITVNGKKATTSEDIIK-- -----NLFNEDTDDAVKRIL LSSCKISEELKKPIKHELN- ---QL---------------  921
R. felis     QKILQDNDI-KITSTFVS-- -----KIFNESLEQTKQRL- RSSNIINAKQYKRIEQYAN- ---KQ---------------  763
R. typhi     AQFTNKNQNT--GS-FL--- ---ENKLFEETREEAILRLL S-LEQLDAQMKKEILSKQGD ILEQLKFINANTEILGSYGQ  243
R. bellii    KDIIRNNKLSPTEKFNLANQ IAANNGIYEKD---DKGNII IDNTNL-------------- --------------------  194

Consensus    ------------ELIRELQG KTNPFEQLEFAYXNAKNLD- --QDIFGNKVDELINNPNIL TIVQQAXFLITEDTNLXK--  968
R. rickettsii ------------ALIRELQG KQNPFEQLEFAYINAKNFD- --QDIFSNRVDELINNPNIL TIVQQANFLITEDTNLRK--  960
R. parkeri   ------------ELIRKLQG KQNPFEQLEFAYINAKNFD- --QDIFGNRVDELINNPNIL TIVQQATFLITEDTNLRK--  959
R. akari     ------------KLMREFES KTTLFEQLEFAYANAKNLD- --QDIFGNKVEELINNPKTL TTAQQATFLITEDTNLRK--  984
R. felis     ------------ECVTEFLR ITNPLEQLKFANKYINILG- --QSTFNGKLNELIENPNKL TFSQKINFVLQGYQELTR--  826
R. typhi     TASTDILKKIPKELPKILSK QGDILEQLKFINANTEILNE HSKAILKDKLKELSKQLDEI SSNQLVGFILDENKINTNLK  323
R. bellii    --------------NNNIIG STSYISNFNPNYKYIISYS- --T------PTEILEKD--- --------LTSEIIDITK--  238

Consensus    --TINXDQAQAKLDDLRTAI LXTIKXEELIX-ANLPQHDF IAIVKEKDPE-LLKEFLKAT TJKX----TGNNNLDQLRLA 1040
R. rickettsii --TINSDQAQAKLDDLRTAI LNTIKFEELIT-TNLPQHDF IAIVKEKDPA-LLQEFLKAT TLKL----TGNNNLDQLRLA 1032
R. parkeri   --TINSDQAQAKLDDLRTAI LSTIKFEELIT-ANLPQHDF IAIVKEKDPE-LLKEFLKAT TLKV----TGNNNLDQLRLA 1031
R. akari     --TIDTDQAQAKLDDLRKAI LSTIKLEELIK-ANLPHNEF IAIVKEKEPE-LLKEFLKAN TIKL----EGNNNLDQLRLV 1056
R. felis     --EIPT--AKANLNKLKQNI LEKIEIQQLIANKDISRKDL LDILNNKNPE-LLKSLLEAK VILEENKLNNSANEVDLKEI  901
R. typhi     NVHFSEKKVRETVSNLNNKI LEKIFLKD---DGTITEQDL TKILQKHQETVLIKNLTKAI VYI-----DGNKNNATVNKT  395
R. bellii    --GFEKDSKGNSYRKFDETK LKQIE-----------QERV YGLIKSKVES---------- --------------------  275

Consensus    LP-SFTXMSNEQJRIL---- ANKLNXXI--ILKALXECSQ EKAK---------------- -------------------- 1077
R. rickettsii LP-SFADMSNEQIRIL---- ANKLKMPI--ILKAIQECSQ EKAK---------------- -------------------- 1069
R. parkeri   LP-SFTGMSNEQIRIL---- SNKLNLSI--ILQALQECSQ EKAT---------------- -------------------- 1068
R. akari     LP-SFTCMSDEQLRVL---- ASKLNMTI--ILNALKEYSQ VKAK---------------- -------------------- 1093
R. felis     IP-SLNYLTSEQLTSL---- INRITIEG--VKTALKAKWQ QENKTVSNNT---------- --------------------  944
R. typhi     LEKCLEQTTPEQQELILDVL THNTRIRTVLITKIEREQRQ NHNKKLNKNIAGDSFVDALK KALVHRTSNSETILKVVEQR  475
R. bellii    -------------------- -------------------- -------------------- --------------------  275

Consensus    ---KXIXTXNMPPPPPPPPX XNSQDLELAYLXSLGITKEX XXXX-----NNNTSKXKTTP KXYXFSSDXALRYKEF---- 1145
R. rickettsii ---QHIHTENMPPPPPPPPL PNAHDLELAYLTSLGITKF- ---------NANTSTFKTTP KTYHFSSDIALRYKEF---- 1132
R. parkeri   ---KYIHTGNMPPPPPPPPL PNSQDLESAYLASLGITKS- ---------TS-TFKTKTTP KTYHFRSDIALKYKEF---- 1130
R. akari     ---KHINTGNMPPPPPPPPS -GLKDAELAYLTTLGITKDW ISRIARLNVKTSTSTFKTTP KIYNFSSDIAVRYKEF---- 1165
R. felis     --EKPLIYNNGTPMPPPIPN GNSNFGTNDYLISMGYTQEF IDRMDKVKPNNNFGKNHN-Y TATDFKSNVGKNYYES---- 1017
R. typhi     KQETPKNLNVWDRISQNIPN LNNQNV------QDENNKEW DESNKNADDLNNTNIYMITK --HDLERAVNETITKFSAMS  547
R. bellii    -------------------- ----LIELAYGANIKYVDYY NSNIND----NSYEKYFTEY E-----KEV-----------  311
```

```
Consensus     -XLSGQKSAG-YKAXYSDA- -------DLLQKAIVEXVAF E-X--------HSKNLSKAX QNN-YFXXI-------ZEAX 1198
R. rickettsii -TLSGQKSAG-YKAKYSDA- -------DLLQKAIVESVAF E----------HSKNLSKAY QNNKYFEQI-------QEAV 1185
R. parkeri    -TLSGQKSAG-YKAAYSDA- -------DLLKKAIVESVAF E----------HSKNLSEAH QNTDYFEQV-------EEAV 1183
R. akari      -ALSGQKSAG-HKAKYSDA- -------DLFQKAIVESVAF E----------HSKNLSKVH QNNTYFAKI-------QEAI 1218
R. felis      ----TSKLGG-TDILLTDS- -------QKLENAIKKEVLA KYIEEPNRDMQDDSLLKQAF EEKFYYAEDKNTKVIPKPSE 1084
R. typhi      TLLKDKKNAGAYQRYLKEAE D-------QL--ALAQEKGK ELIKNSA---QTFKIIPKKY QDD-----I-------NENW  603
R. bellii     ----ITKALG-YKRAYEEKT GLYKSEVDKLDRSILSEE-- EYIKKCE------EILEK-- --------------IGESSW  362

Consensus     DTMHXSFIGPRTEIGQEIHN -------------------- -----------IYT----SK LLXL--TKDKEFIKYVEDNI 1241
R. rickettsii DTMHPRFIGPRTEIGQNIHN -------------------- -----------IYT----SK LLEL--TKDKEFIKYVED-I 1227
R. parkeri    NTMHSSFTGPRTEMGQIIHN -------------------- -----------IYI----SK LLEL--TKDKEFIKYVEDNI 1226
R. akari      DTMHSSFIGPRTEIGQEIHN -------------------- -----------IYT----SK LLAL--TKDKEFIKYVEDDI 1261
R. felis      VNFDPNFIGPRTEVGQEIYE -------------------- -----------LYE----QE LLKL--ARDPVFIEYVKNNN 1127
R. typhi      QN----YLSPAEMIELTALN EHTNTLKSNKNKSGHFRSSE EALQYKAKQHEYYTLLAELK KIGIAKQQEKLVKDYVDEMI  679
R. bellii     DTYQVSKPGSSSKISYEEYE ----KLVTNEAKKR-----A VIARFLMEDNNKIN----GK RLKI--VKEDGSEEYIDNTE  427

Consensus     ILSKKLTEAFTSAXS----- -------------------- -------------------- -------------XFIGPRT 1263
R. rickettsii ILSKKLTEAFTSADS----- -------------------- -------------------- -------------DFIGPRT 1249
R. parkeri    ILNKKLTKAFTSAGS----- -------------------- -------------------- -------------GFIDSRT 1248
R. akari      ILSKKLTEAFTSADS----- -------------------- -------------------- -------------DFIGPRT 1283
R. felis      NTQ----------------- -------------------- -------------------- -------------------- 1130
R. typhi      TNAKQAVEKFERTSLEHINQ KKENKQISKEILDAQERLEN AKQKIEFIKFKYIISNKRQV NSSDEDSDDDADKNAIKQKT  759
R. bellii     FN------------------ -------------------- -------------------- --------------LIRAEN  435

Consensus     EXXQXXHDIY------IQQL AKYPEE-XVKEAFNTXXPDF IGPRTEIGQE---VHNIYKS XLLELXKDKELFLFXEQJLA 1333
R. rickettsii EIGQKIHDIY------IQQL TKYPEE-AVKEAFNTAHPDF IGPRTEIGQE---VHNIYKS QLLELAKDKELFLFVQQVLV 1319
R. parkeri    RLEQKIHDIY------IQQL TKYPEEEAVKEAFNTANPDF IGPRTEIGQE---VHNIYKS QLLELTKDQELFLFTQQILA 1319
R. akari      EIGQEVHNIY------TQQL AKYPEE-TVKEAFNTANSDF IGPRTEIGQE---VHNIYKS KLLELAKDKELFLCVEQLLA 1353
R. felis      -------------------- -------------------- -------------------- ----KDERELLISFIEQIES 1146
R. typhi      ELENAQKDINQAKKNLEDAK AKYAAL---QITLNY-SPNG MDSKTTEDQEQFKTDNIAAE S---LEDMAVMFKDSEEAAE  832
R. bellii     --------NW------DLTP ANIPLSKVMK---------- ---------D---TNN---- -LSENEKMKKLLARIEFLEN  474

Consensus     ESAELEQKYGS---DIQSEN XNQEKKVG------------ ----------------L--- -------------------- 1359
R. rickettsii DSAELEQKYGS---DIQSEN SNDEKKVG------------ -------------------- -------------------- 1344
R. parkeri    ESTELEQKYGS---DIQSEN SDNEKTVGR----------- ----------------LDNE KTVGRLDNEKTVGRLDNEKT 1369
R. akari      ESAELEQKYGT---DLQSEN NNQEKKVG------------ -------------------- -------------------- 1378
R. felis      KRPELEQKYGS---DVQSED NNQEKKVGH----------- ----------------LNM- -------------------- 1175
R. typhi      RKEEVTLNYSPNGMDSKTTE DQEQFKTDNIAAESLEDMAV MFKDSEEAAERKEEVTLNYS PNGMDSKTTEDQEQFKTDNI  912
R. bellii     NTNNL---------DIINEL EDAKRQLDE----------- ----------------L--- -------------------- 495

Consensus     --RLDKKKLRLFXQENEAAN -------------------- DESSTKDDTQ-PED------ SNKKSEZS-----DSXTALS 1405
R. rickettsii --RLDKTKLRLFQQENEATN -------------------- DESSTKDDTQ-PED------ SNKKSEQS-----DSKTALS 1390
R. parkeri    VGRLDQEKLQSFKQENEATN -------------------- DESSTKDDTQ-PED------ SNKKSEQS-----DSKTALS 1417
R. akari      --RLDPKQLRLFLQANESAN -------------------- DESSTKDNTQ-PED------ SNKKSEEF-----DSETALS 1424
R. felis      -----KQFQSLFQQENESAN -------------------- DESSTKDDPQ-PED------ SNKKSEKS-----DSETALS 1218
R. typhi      AAESLEDMAVMFKDSEEAAE RKEEVTLDYSPNGMDSKTTE DQEQFKTDNIAAESLEDMAV MFKDSEEAAERKEEVTLDYS  992
R. bellii     ---KSKKITGLFALNNNGEN -------------------- ASISFDQDLIDILK------ ILKEVPDF-----VSIIRAH  541

Consensus     PRLLSSNDSKNDKS--SD-- ----DKKSL-----X----- -------------------- -------------------- 1427
R. rickettsii PRLLSSNDSKNDKS--SD-- ----YKKSLLELRSSDED-- -----D-----QGYATGYTT DEEELEESNSTTGEELKKDI 1450
R. parkeri    PRLLSSNDSKNDKS--SD-- ----DKKSL----------- -------------------- -------------------- 1438
R. akari      PRLLSSNDSKNDKS--SD-- ----N--------------- -------------------- -------------------- 1441
R. felis      PRLLSSNDSKNDKS--SD-- ----DKKSLLVLRSSE---- -------------------- -------------------- 1246
R. typhi      PNGMDSKTTEDQEQFKTDNI AAESLEDMAVMFKDSEEAAE RKEEVTLNYSPNGMDSKTTE DQEQFKTDN-IAAESLED-- 1069
R. bellii     PDLFPTV------------- -------------------- -------------------- --------------------  548

Consensus     -------XALRSSDEDDX-- -------------------- GYXTDEXEL------EE-XN STT-EEXKKDIALESEDEAI 1470
R. rickettsii SDYKKSLLALRSSDEDDQ-- -------------------- GYATDEEEL------EE-GN STTGEELKKDIVLESEDEAI 1501
R. parkeri    -------LALRSSDEDDT-- -------------------- GYATDEEEL------EE-SN STTDEELKKDVVLESEDEAI 1482
R. akari      -------IDLRSSDEEDK-- -------------------- GYETDE-EL------KE-SN NTTKEESQKDIALESEDEAI 1484
R. felis      -------------------- -------------------- -------------------- ----EESKKDIALESEDEAI 1262
R. typhi      -------MAVMFKDSEEAAE QKEEVNRQHHEEQNRQKQEH NCLDTEEEVVHKEKIADLTA ETETKVFKKEIAL-EENEAM 1141
R. bellii     -----------FNDPDTL-- -------------------- AFL------------E---- KCS-PE-------DYENIII  571
```

| | | | | | |
|---|---|---|---|---|---|
| Consensus | DVSFKTEAIXEQDEATQRQQ | VSDDTSRKVAILVXATSTLH | KPVHYNILSDR-LKVAAIGA | GDEETSINRGVWISGLYGIN | 1549 |
| *R. rickettsii* | DVSFKTEAITEQDKVTQRQQ | VSDDTSGKVAILVQATSTLH | KPVHYNI-NDR-LTVAAIGA | GDEETSINRGVWISGLYGIN | 1579 |
| *R. parkeri* | DVSFKTEAITEQDEVTQRQQ | VSDYTSGKVAILVQATSTLH | KPVHYNI-NDR-LTIAAIGA | GDEETSINRGVWISGLYGIN | 1560 |
| *R. akari* | DVSFTTEAIAEQDEATQRQQ | VSDDTSRKVAILAKATSTLH | KPVHYNILSDR-LKVSAIGA | GDEEPIIARGVWISGLYGMN | 1563 |
| *R. felis* | DMSFKTEAIAEQDEATQRQQ | VSDDTNRKVAILVKATSTLH | KPVHYNILSDR-LKVAAIGA | GDEEASINRGVWISGLYGIN | 1341 |
| *R. typhi* | DVSFKTETIVEQDEAIQRQQ | VSDDTSRKVAILVKATSTLH | KPVHHNILSDR-LKVTVIGA | GDEKTNINRGLWISGLYGVN | 1220 |
| *R. bellii* | ELSLLDSKSQDLESL----I | VKKQISEEVTLHNQVASITN | KPIHMGIHGRLLLPTAAITG | GDEEDAINRGVWISGLYGVS | 647 |

| | | | | | |
|---|---|---|---|---|---|
| Consensus | KQGXWKNIPKYQGRTTGITI | GADAEFINSHDVIGIAYSRL | ESQIKYNKKLGKTAVNGHLL | SIYGLKELIKGFSLQXITSY | 1629 |
| *R. rickettsii* | KQSIWKNIPKYQNRTTGITI | GADAEFINSHDVIGIAYSRL | ASQIKYNKKLGKTAVNGHLL | SIYSLKELIKGFSLQTITSY | 1659 |
| *R. parkeri* | KQRIWKNIPKYQNRTTGITI | GTDAEFINSHDVIGIAYSRL | ESQIKYNKKLGKTTVNGHLL | SIYSLKELINGFSLQTITSY | 1640 |
| *R. akari* | KQGTWKNIPKYQGRTTGITI | GADVEFINSHDVIGIAYSRL | ESKIKYNKKLGTIAVNGHLF | SIYGLKELIKDLSLHAITSY | 1643 |
| *R. felis* | KQGTWKNIPKYQGRTTGVTI | GADAEFINSHDVIGIAYSRL | EFQIKYNKKLEKTAVNGHLL | SIYGLKELIKGFSLQAITSY | 1421 |
| *R. typhi* | KQGSWKNIPKYQGRTTGFTI | GADAEFINNHDVIGIAYSNL | ESKIKYNKKLGKTAVYGHLL | SIYGLKELIPGFSLHTIVSY | 1300 |
| *R. bellii* | NQKAWRSIPKYQGRTSGVTI | GMDTELSNSSDVIGIAYSRI | ESHFKYNKKFAKTALNGHLL | SVYGLKELPKNFSLQAIASV | 727 |

| | | | | | |
|---|---|---|---|---|---|
| Consensus | GHNYIKNKSKNINNIIGKYQ | NNNLSFQTLLNYKYRTKYDL | HFIPNIGFKYDYSRASNYKE | YNVDIENLMIQKKSNQLFES | 1709 |
| *R. rickettsii* | GHNYIKNRSKNINNIIGKYQ | NNNLSVQTLLNYKYRTKYDL | HFIPNIGFQYDYSRASNYKE | YNVDIENLMIQKKSNQLFES | 1739 |
| *R. parkeri* | GHNYIKNRSKNINNIIGKYQ | NNNLSFQTLLNYKYRTKYDL | HFIPNIGFQYDYSRASNYKE | YNVDIENLMIQKKSNQLFES | 1720 |
| *R. akari* | GHNYVKNKSKNINNIIGKYQ | NNNLSFESLLNYKYHTKYDL | HFIPNIGFKYDYSRASNYKE | YNVDIENLMIQKKSNQLFES | 1723 |
| *R. felis* | GHNYIKNKSKSINNIIGKYQ | NNNLSFQTLLNYKYRTKYDL | HFIPNIGFKYDYSRASNYKE | YNVDIENLMNQKKSNQSFES | 1501 |
| *R. typhi* | GYNYIKNKSKNLDKIIGKYQ | NNNLSFQTLLNYKYCTKYNL | HFIPSIGFKYDYSRASNYKE | YNIDIENLMIQKKSNQSFES | 1380 |
| *R. bellii* | GHNYIKNKATTANNIIGKYQ | NNNFNFEALLNYKYRINYNL | YLIPNIGLKYDYSRSSGYKG | NNF-VQKLMIQKKSNRLLTT | 806 |

| | | | | | |
|---|---|---|---|---|---|
| Consensus | SJGGKIVFKPIXTTNNIVLT | PSLYGNIERHFNNKNTKVNA | KATFKGQTLQETIIIPKQPK | LGYNIGSNILMSRKNINVLL | 1789 |
| *R. rickettsii* | SLGGKIVFKPIVTTNNIVLT | PSLYGNIEHHFNNKNTKVNA | KATFKGQTLQETIITLKQPK | LGYNIGSNILMSRKNINVLL | 1819 |
| *R. parkeri* | SLGGKIVFKPIVTTNNIVLT | PSLYGNIEHHFNNKNTKVNA | KATFKGQTLQETIITLKQPK | LGYNIGSNILMSRKNINVLL | 1800 |
| *R. akari* | SIGGKIVFKPIATISNVVLT | PSLYGNIERHFNHKNTKVNA | KATFKGQILQETIIIPKQPK | LGYNIGSNMLMSRKNINVLL | 1803 |
| *R. felis* | SIGGKIVFKPIATVNNIILT | PSLYGNIERHFNNKNTKVNA | KATFKGQTLQETIIIPKQPK | LGYNIGSNILMSKKNINVLL | 1581 |
| *R. typhi* | SIGGKIVSKPIIITNNTILT | LSVHGNIERHFNNKNTKVNA | KATFKKQTLQETIIIPKQPK | FGYNIGNNILMSIKNINVLY | 1460 |
| *R. bellii* | SLGSKVEFKSIKVLNDITLV | PSLYGSIENHFYNKDTKVNA | KAVLNNQIIEEKIIISKQPK | FGYNIGGNVLLTKKNINVVF | 886 |

| | | | |
|---|---|---|---|
| Consensus | EYNYYTHRKYQSHQGLIKLK | VNL | 1812 |
| *R. rickettsii* | EYNYYTHRKYQSHQGLMKLK | VNL | 1842 |
| *R. parkeri* | EYNYYTHRKYQSHQGLIKLK | VNL | 1823 |
| *R. akari* | EYNYYTHRKYQSHQGLVKLK | VNL | 1826 |
| *R. felis* | EYNYYTHRKYQSHQGLIKLK | VNL | 1604 |
| *R. typhi* | EYNYYTHKKYHSHQGLIKLK | INL | 1483 |
| *R. bellii* | EYNHYTHKKYKSHQGLVKLK | INL | 909 |

**FIGURE S1.** Sequence alignments of RickA and Sca2 orthologs from divergent *Rickettsia* species. (A) Geneious sequence alignment of *Rickettsia* RickA proteins. Spotted fever group II (*R. akari* and *R. felis*) RickA proteins have two WH2 domains. (B) Clustal Omega sequence alignment of *Rickettsia* Sca2 protein orthologs. The domains are indicated with colored text in the alignment according to the same scheme as FIGURE 1. Selected species are part of the following groups: SFG I = *R. rickettsii* and *R. parkeri*, SFG II = *R. akari* and *R. felis*, TG = *R. typhi*, BG = *R. bellii*.

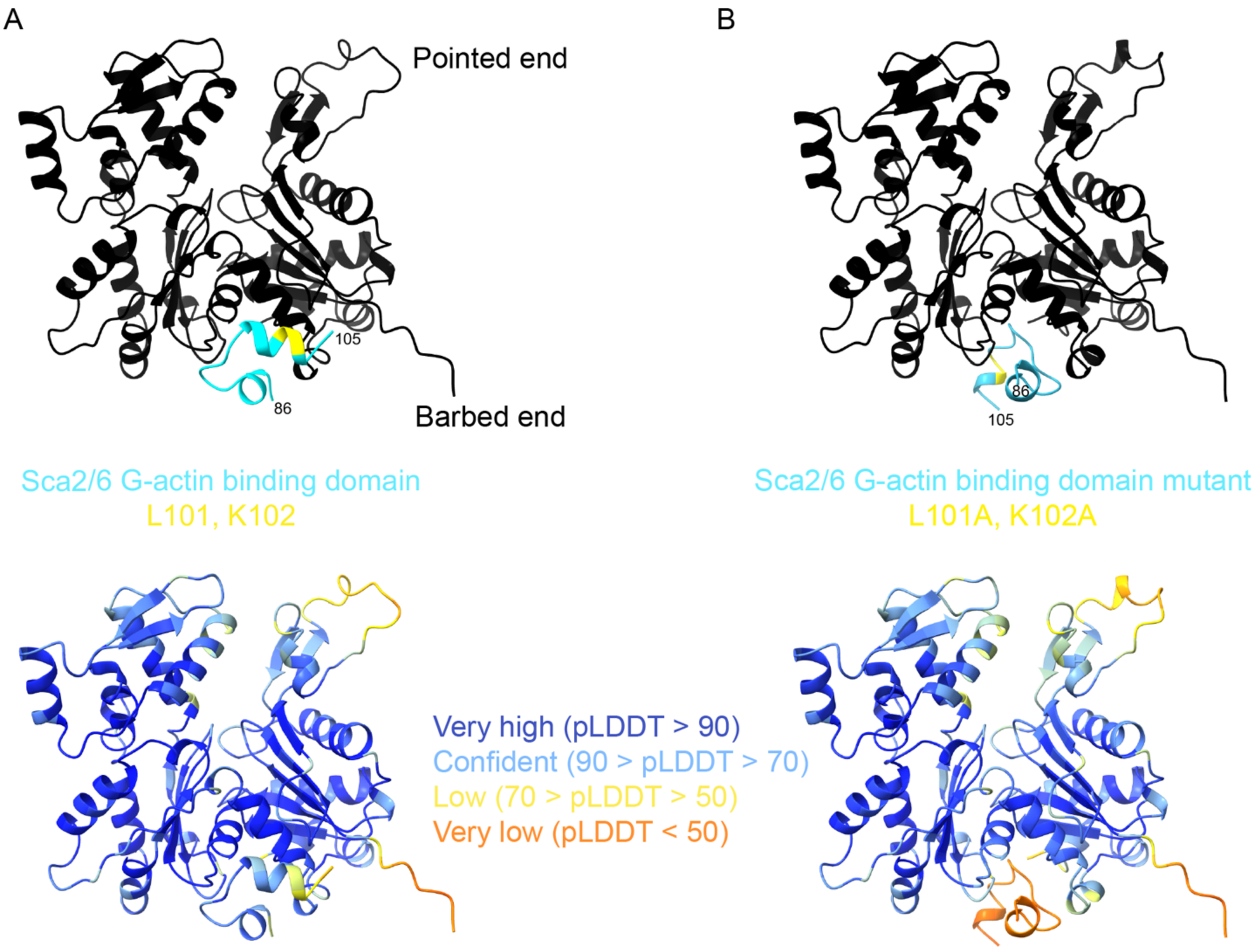

**FIGURE S2.** The NTD of *R. bellii* Sca2/6 is predicted to bind G-actin. (A) AlphaFold3 prediction of *R. bellii* Sca2/6 G-actin binding domain (light blue, aa 86 – 105) bound to the barbed end of a rabbit skeletal muscle actin monomer (black, UniProt P68135). L101 and K102 are highlighted (yellow). The same AlphaFold3 prediction is color coded below by pLDDT confidence values. (B) R. bellii Sca2/6 aa 86-105 including the L101A and K102A mutations bound to rabbit skeletal muscle actin, color coded as in (A).

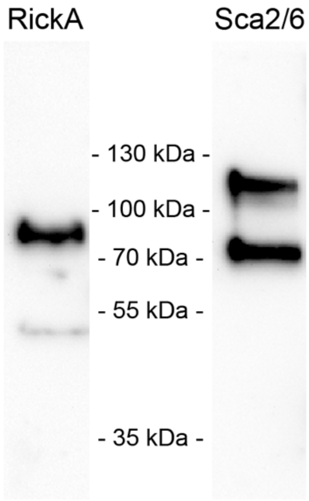

**FIGURE S3.** RickA and Sca2/6 are expressed in *R. bellii*. Immunoblot of wild type *R. bellii* 30% preparation stock probed with rabbit anti-RickA or anti-Sca2/6 antibodies.

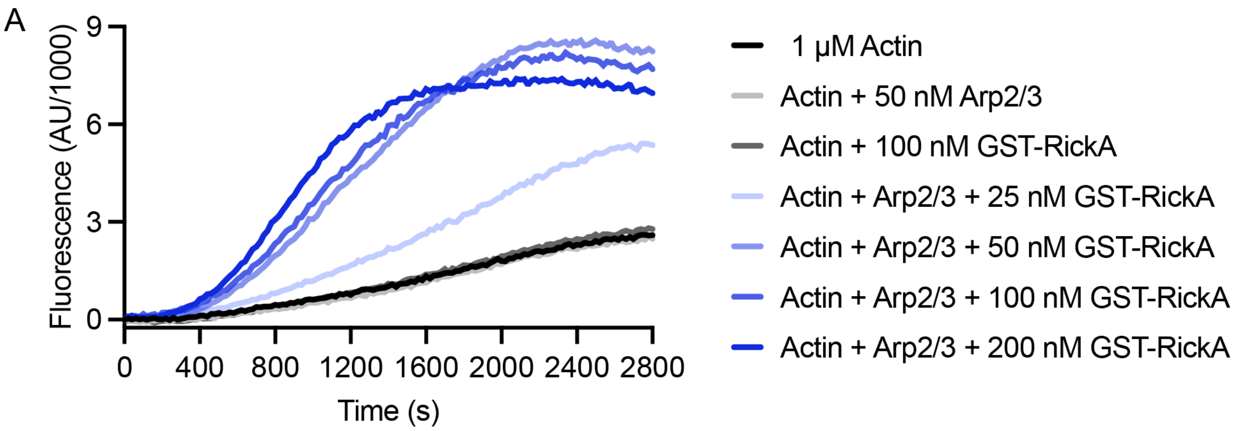

**FIGURE S4.** *R. bellii* RickA nucleates actin filaments through the activation of the host Arp2/3 complex. Kinetics of assembly of 1 μM actin (10% pyrene-labeled) performed with or without 50 nM Arp2/3 complex and the indicated concentrations of *R. bellii* GST-RickA.

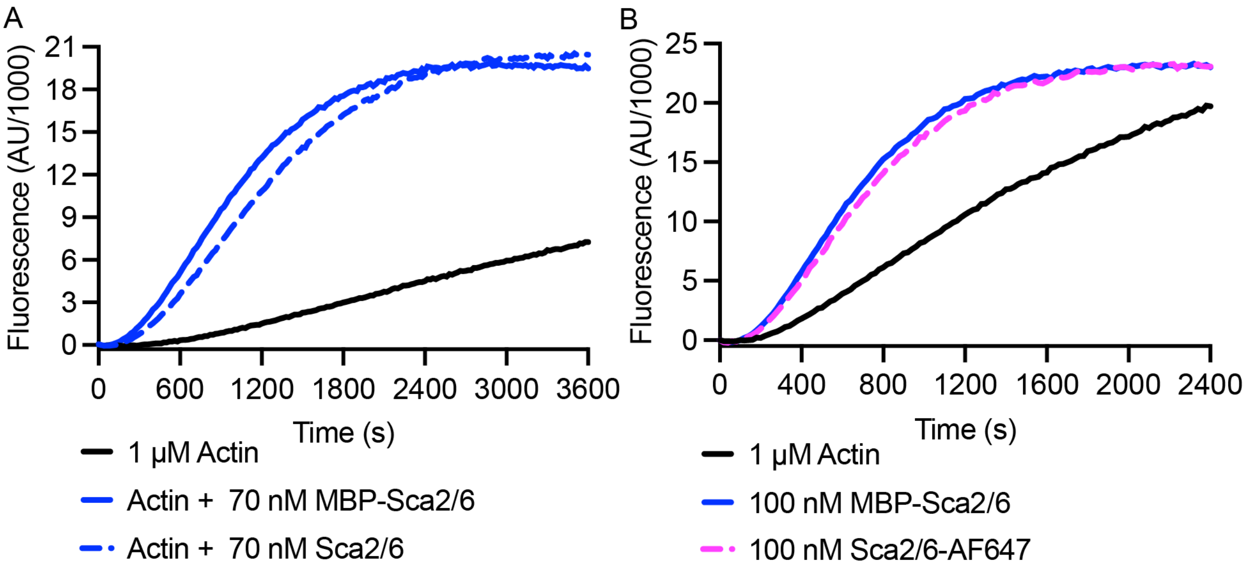

**FIGURE S5.** Cleaved *R. bellii* Sca2/6 and maleimide labeled Sca2/6-AF647 actin nucleation activity is comparable to that of MBP-Sca2/6. (A) and (B) Kinetics of assembly of 1 μM actin (10% pyrene-labeled) performed with the indicated versions of *R. bellii* Sca2/6.

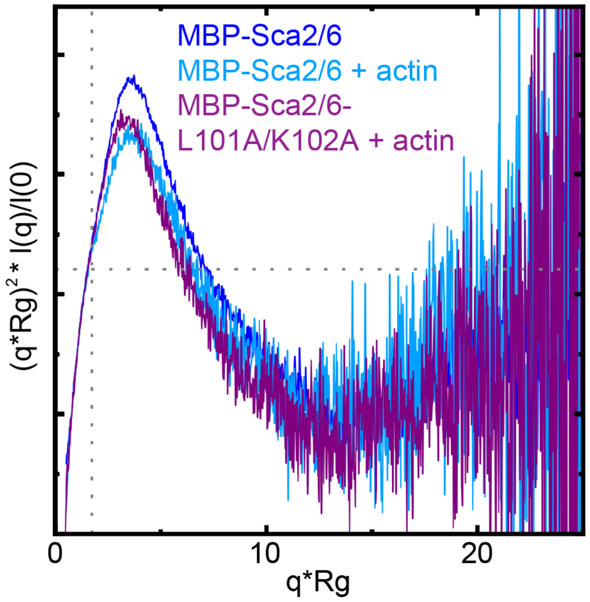

**FIGURE S6.** Experimental SAXS curves for MBP-Sca2/6, MBP-Sca2/6 plus G-actin, and MBP-Sca2/6-L101A/K102A plus G-actin are shown as normalized Kratky plots. Dashed lines indicate where a globular protein would peak.

**MOVIE S1.** *R. bellii* Sca2/6 does not track the growing ends of actin filaments. TIRF elongation assay performed with 1 μM actin (33% rhodamine-labeled) and 0.5 or 5 nM *R. bellii* Sca2/6-AF647 imaged every 5 s. Movie is shown at 7 frames per s. Scale bar = 5 μm.

**MOVIE S2.** *R. parkeri* actin-based motility is rapid and directed. Live confocal imaging of *R. parkeri* (blue) with actin tails (magenta) in A549 human epithelial cells and HMEC-1 human endothelial cells at 24 hpi. Images were acquired every 5 s. Movie is shown at 7 frames per s. Scale bar = 5 μM.

**MOVIE S3.** *R. belli* actin-based motility is less rapid and more meandering. Live confocal imaging of *R. bellii* (blue) with actin tails (magenta) in A549 human epithelial cells and HMEC-1 human endothelial cells at 24 hpi. Images were acquired every 5 s. Movie is shown at 7 frames per s. Scale bar = 5 μM.
